# Loss of Tumour Suppressor TMEM127 Drives RET-mediated Transformation Through Disrupted Membrane Dynamics

**DOI:** 10.1101/2023.06.28.546955

**Authors:** Timothy J. Walker, Eduardo Reyes-Alvarez, Brandy D. Hyndman, Michael G. Sugiyama, Larissa C.B. Oliveira, Aisha N. Rekab, Mathieu J.F. Crupi, Rebecca Cabral-Dias, Qianjin Guo, Patricia L.M. Dahia, Douglas S. Richardson, Costin N. Antonescu, Lois M. Mulligan

**Author notes:** To whom correspondence should be addressed:* Dr. Lois M. Mulligan, Division of Cancer Biology and Genetics Cancer Research Institute at Queen’s University Botterell Hall Rm A315, Queen’s University Kingston, ON, Canada K7L 3N6.

## Abstract

Internalization from the cell membrane and endosomal trafficking of receptor tyrosine kinases (RTK) are important regulators of signaling in normal cells that can frequently be disrupted in cancer. The adrenal tumour pheochromocytoma (PCC) can be caused by activating mutations of the RET receptor tyrosine kinase, or inactivation of TMEM127, a transmembrane tumour suppressor implicated in trafficking of endosomal cargos. However, the role of aberrant receptor trafficking in PCC is not well understood. Here, we show that loss of TMEM127 causes wildtype RET protein accumulation on the cell surface, where increased receptor density facilitates constitutive ligand-independent activity and downstream signaling, driving cell proliferation. Loss of TMEM127 altered normal cell membrane organization and recruitment and stabilization of membrane protein complexes, impaired assembly, and maturation of clathrin coated pits, and reduced internalization and degradation of cell surface RET. In addition to RTKs, TMEM127 depletion also promoted surface accumulation of several other transmembrane proteins, suggesting it may cause global defects in surface protein activity and function. Together, our data identify TMEM127 as an important determinant of membrane organization including membrane protein diffusability, and protein complex assembly and provide a novel paradigm for oncogenesis in PCC where altered membrane dynamics promotes cell surface accumulation and constitutive activity of growth factor receptors to drive aberrant signaling and promote transformation.

## Introduction

Internalization of activated receptor tyrosine kinases (RTK) from the cell membrane and trafficking through endosomal compartments is a critical mechanism regulating the magnitude and duration of downstream signals (Schmid, 2017). RTKs internalized by clathrin mediated endocytosis (CME) transition through successive early and late endosomes for eventual degradation in lysosomes (Sorkin and Fortian, 2015). In cells where endocytosis or trafficking is impaired, RTKs may accumulate inappropriately at the cell surface or in aberrant subcellular compartments, contributing to cell transformation and oncogenic growth. Despite the importance of CME and RTK cargo transition through compartments, the trafficking proteins required for transitions and their roles in these processes remain poorly understood.

RET (Rearranged during Transfection) is a receptor tyrosine kinase required for development of the kidneys, nerves and neuroendocrine cell types (Mulligan, 2019). Under normal conditions, RET is activated by binding of its glial cell line-derived neurotrophic factor (GDNF) ligands and GPI-linked GDNF family receptor α (GFRα) coreceptors, which together recruit RET complexes to lipid rafts, leading to downstream signalling through multiple pathways (Mulligan, 2019; Pierchala et al., 2006).

Activated RET at the cell membrane is internalized via CME to early endosomes where signaling persists, and subsequently trafficked through the endolysosomal system for down regulation and degradation (Crupi et al., 2015; Richardson et al., 2012). Activating mutations of RET or over expression of the wildtype (WT) protein are established drivers in several cancers (Mulligan, 2019). Constitutively active RET mutants give rise to the inherited cancer syndrome multiple endocrine neoplasia type 2 (MEN2), which is characterized by the adrenal tumour pheochromocytoma (PCC), while activating mutations or increased RET expression are found in a subset of sporadic PCC (Le Hir et al., 2000; Mulligan, 2019; Takaya et al., 1996a; Takaya et al., 1996b).

PCC has also been associated with inactivating mutations of TMEM127, a multipass integral membrane protein broadly expressed at low levels and localized to the plasma membrane, endosomes and lysosomes in normal cells (Deng et al., 2018; Qin et al., 2014; Qin et al., 2010). TMEM127 functions are poorly understood but it has been suggested to facilitate endosomal transition and cargo trafficking and has been shown to modulate endolysosomal function in renal cell carcinomas (Deng et al., 2018; Qin et al., 2010). Loss of function mutations of *TMEM127* have been identified in both familial and sporadic PCC, leading to loss of TMEM127 protein or mislocalization to the cytosol in tumour cells (Neumann et al., 2019; Qin et al., 2010). Interestingly, our recent data have demonstrated that RET expression is increased in TMEM127 mutant PCC (Guo et al., 2023)

The presentation of RET mutant or TMEM127 deficient PCC are not clinically distinct. Both RET and TMEM127 have been predicted to confer PCC susceptibility through altered kinase signaling (Mulligan, 2019; Neumann et al., 2019; Qin et al., 2010), however, their relationship has not been explored. The importance of endolysosomal trafficking in regulating the intensity and duration of RET pro-proliferative signals, suggested that loss of TMEM127-mediated endosomal transition could alter RET trafficking, promoting accumulation or signaling from aberrant subcellular compartments and contributing to transformation. In this study, we have investigated the effects of TMEM127 depletion on RET regulation and function and more broadly on cellular processes that could contribute to PCC pathogenesis. Here, we show that loss of TMEM127 leads to cell surface accumulation and constitutive activation of RET and that this accumulation is due to alterations in membrane organization and composition that reduce efficiency of CME and decrease internalization of RET as well as other transmembrane proteins, suggesting a global impairment of membrane trafficking. As a result of changes in membrane dynamics, TMEM127 depletion increased RET half-life and led to constitutive RET-mediated signalling and cell proliferation that was blocked by RET inhibition. Our data further demonstrate that TMEM127 loss increases membrane protein diffusability and impairs normal membrane transitions and the stabilization and assembly of membrane protein complexes, allowing inappropriate accumulation of actively signaling RET molecules at the cell membrane, and that mis-localized RET is the pathogenic mechanism in TMEM127-mutant PCC.

## Results

### Loss-of-function mutations disrupt TMEM127 localization and reduce early endosome formation

WT-TMEM127 has previously been shown to localize to endosomal compartments, while loss-of-function PCC-TMEM127 mutants reduce protein expression or may be aberrantly localized in the cytosol (Flores et al., 2020; Qin et al., 2010). Here, we assessed colocalization of transiently expressed TMEM127 with the early endosome marker EEA1 in human retinal pigment epithelial ARPE-19 cells by immunofluorescence microscopy. We showed that WT-TMEM127 localized robustly to EEA1-positive endosomes but that colocalization was significantly reduced for two transmembrane domain TMEM127 mutants (C140Y, S147del), which did not show appreciable localization to EEA1 puncta (Figure 1A, B). Interestingly, the total numbers of EEA1 puncta in cells were modestly reduced in the presence of TMEM127-S147del and significantly reduced for TMEM127-C140Y (Figure 1B), suggesting a systemic disruption of endosomal formation or stability in response to loss of TMEM127 function in these cells. In ARPE-19 cells transiently expressing RET and TMEM127, GDNF stimulation promoted colocalization of RET with WT but not mutant TMEM127 in EEA1 positive endosomes (Figure 1C). RET and EEA1 colocalization was reduced but not eliminated in the presence of TMEM127 mutants, suggesting TMEM127 loss may limit RET localization to early endosome compartments.

**Figure 1.**
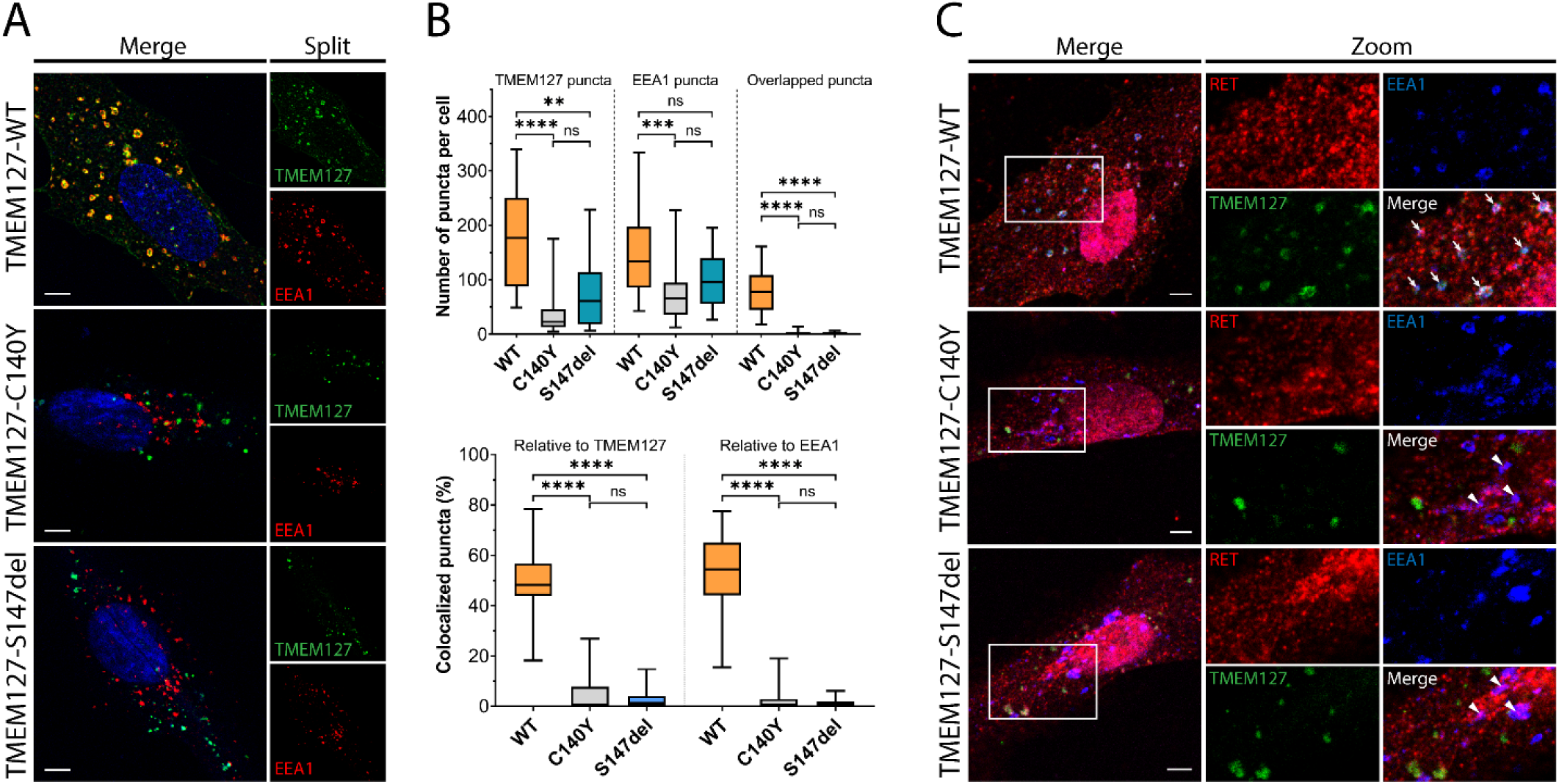
Wildtype and mutant TMEM127 are differentially localized in endosomes. (**A**) Immunofluorescence confocal images of ARPE-19 cells transiently expressing EGFP-TMEM127-WT, -C140Y, or -S147del, and stained for early endosome marker EEA1 (red) and Hoechst nuclear stain (blue). Yellow puncta in the merged images indicate EEA1 colocalization with WT but not mutant TMEM127. (**B**) Quantification of puncta in cells expressing the indicated TMEM127 proteins under conditions in **A**. Quantification of TMEM127 positive or EEA1 positive and overlapping puncta (upper panel) and percent colocalized puncta relative to TMEM127 or EEA1 (bottom panel). Data from two independent experiments representing 17-31 cells per condition are shown (Kruskal-Wallis test and Dunn’s multiple comparisons test; **p<0.01, ***p<0.001, ****p<0.0001). Scale bars = 5 μm. **C**) Immunofluorescence confocal images of ARPE-19 cells transiently expressing RET, GFRα1 and EGFP-TMEM127-WT, -C140Y, or -S147del, treated with GDNF (100 ng/ml) for 10 minutes and stained for RET (red) and EEA1 (blue). White puncta (white arrows) indicate RET, TMEM127-WT, and EEA1 colocalization. Colocalization of RET and EEA1 but not TMEM127 in cells expressing TMEM127-C140Y or TMEM127-S147del mutants is indicated with white triangles. Representative images from 14-25 cells per condition. Scale bars = 5 μm. Zoom Box = 15 μm x 22 μm.

### TMEM127 depletion promotes cell surface RET accumulation

Human PCC cell lines and patient derived organoid cultures have not been successfully generated to date. We therefore generated a CRISPR-Cas9 TMEM127-knockout (KO) and control (Mock-KO) in human neuroblastoma cell line SH-SY5Y, which endogenously expresses RET and TMEM127 (Figure 2A), to explore the effects of TMEM127 loss of function on RET localization and functions. We confirmed that these TMEM127-KO cells had similarly reduced EEA1 positive endosomes to our transient models, described above (Figure 2-figure supplement 1A). Interestingly, we observed notably increased levels of RET protein in TMEM127-KO compared to control Mock-KO cells (Figure 2A), which was consistent with accumulation of a fully mature glycosylated RET form that is found primarily on the cell surface (Richardson et al., 2006; Richardson et al., 2012; Takahashi et al., 1991). This accumulation was not due to changes in *RET* gene transcription, since TMEM127-KO cells had significantly less RET mRNA in qRT-PCR assays (Figure 2-figure supplement 1B), suggesting a possible feedback loop modulating RET transcription. Transient re-expression of WT-TMEM127 in these cells reduced RET protein levels (Figure 2-figure supplement 1C). Using a surface protein biotinylation assay (Figure 2A, Figure 2-figure supplement 2, (Reyes-Alvarez et al., 2022)) to specifically detect plasma membrane proteins, we showed that cell surface RET was significantly increased, by approximately 5 fold, in TMEM127-KO compared to Mock-KO SH-SY5Y cells. We also saw similar plasma membrane accumulation of endogenous N-cadherin and transferrin receptor-1 proteins in HEK293 cells depleted for TMEM127, and reintroduction of flag-tagged TMEM127 significantly reduced membrane localization (Figure 2-figure supplement 1D). Taken together, the accumulation of RET on the cell surface and reduced RET in early endosomes suggested that TMEM127 loss might disrupt RET internalization.

**Figure 2.**
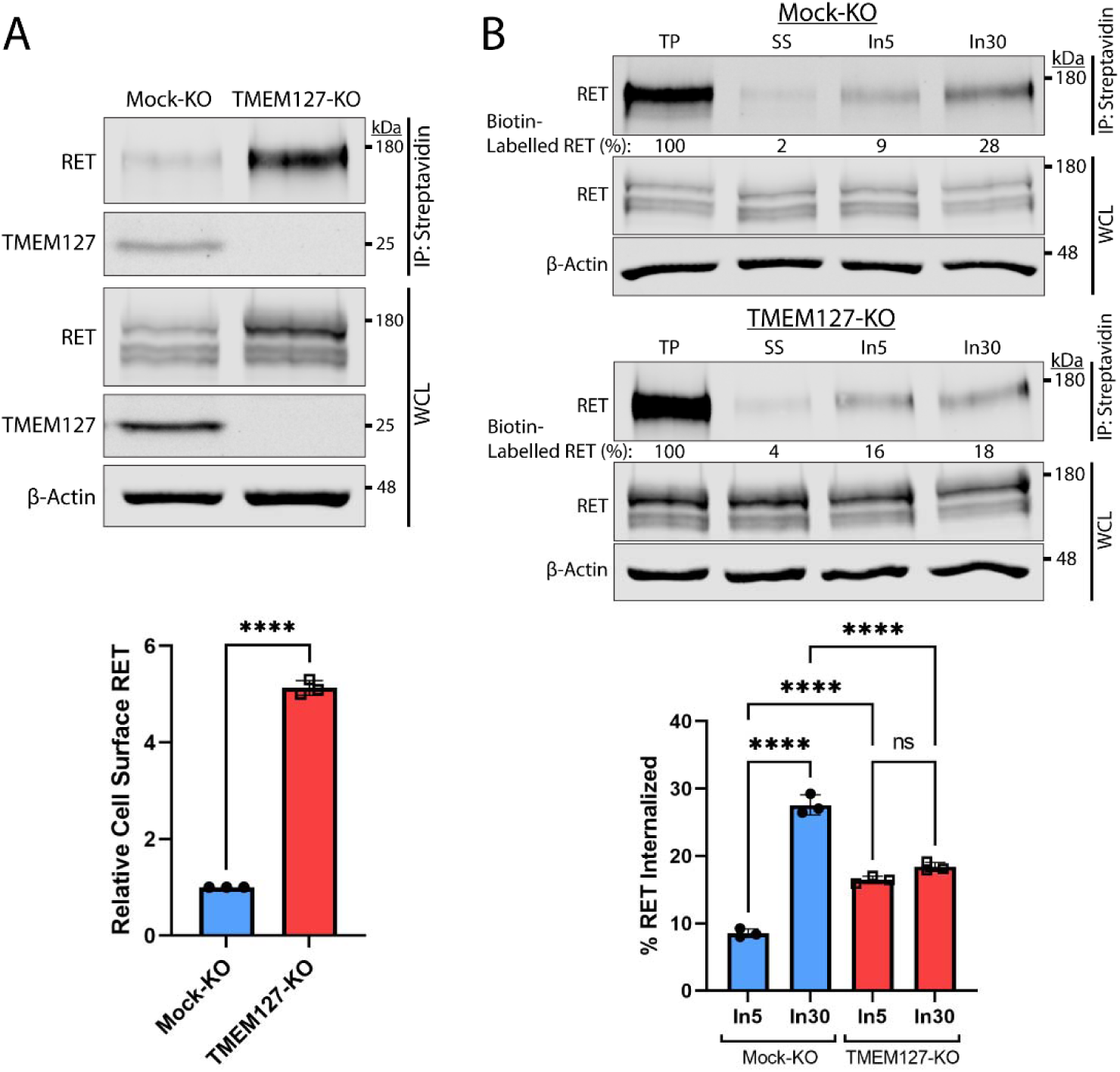
RET accumulates on the cell surface due to reduced internalization in the absence of TMEM127. (**A**) Immunoblot showing total and cell surface biotinylated RET and TMEM127 proteins in Mock-KO and TMEM127-KO SH-SY5Y cells. Surface proteins were biotinylated and collected by streptavidin immunoprecipitation (IP: Streptavidin). Whole cell lysate (WCL) and IP samples were separated by SDS-PAGE and immunoblotted for RET and TMEM127 (upper panel). β-Actin was used as a loading control. Biotinylated cell surface RET levels (IP) were normalized to total RET (WCL) and expressed relative to Mock-KO (bottom panel). Fold increase in relative surface RET is indicated for TMEM-KO cells. Three independent experiments (n=3) are shown as mean ±SD (Two-tailed unpaired t-test; ****p<0.0001). (**B**) Immunoblots showing internalization of biotinylated cell-surface RET protein in Mock-KO and TMEM127-KO SH-SY5Y cells (top). Surface proteins were biotin labelled (TP; total protein) and either cell-surface biotin stripped (SS; Surface Strip control) or incubated at 37°C with GDNF (100 ng/ml) for 5 (In5) or 30 (In30) min to allow internalization, and remaining cell-surface biotin was stripped. Biotinylated proteins were collected, separated, and immunoblotted as in **A**. β-Actin was used as a loading control. Mean ±SD of internalized biotinylated RET, relative to TP of three independent experiments (n=3) are shown in the bottom panel (One-way ANOVA and Tukey’s multiple comparisons test; ****p<0.0001).

### TMEM127 depletion promotes RET accumulation through impaired internalization

We assessed RET internalization in response to GDNF over time in TMEM127-KO and Mock-KO cells using a biotinylation internalization assay (Figure 2B, Figure 2-figure supplement 2, (Reyes-Alvarez et al., 2022)). Consistent with our previous studies (Crupi et al., 2015; Richardson et al., 2012), minimal RET internalization was detected by 5 min of GDNF treatment in Mock-KO cells and increased over time, with significantly greater internalization detected by 30 min (Figure 2B). In TMEM127-KO cells, RET internalization was similar at all time points and did not increase over time with longer GDNF treatments, with significantly less internalization observed after 30 min compared to Mock-KO controls (Figure 2B). Recycling of internalized RET to the cell surface in biotinylation assays was also reduced in TMEM127-KO compared to Mock-KO cells with a smaller proportion of the internalized RET being returned to the cell surface (Figure 2-figure supplement 2) suggesting RET surface accumulation was not due to enhanced recycling. Our data show that RET internalization into TMEM127-KO cells is not GDNF responsive and remains at a low constitutive rate despite high levels of RET protein accumulating on the cell surface, suggesting that TMEM127 depletion may impair cellular internalization mechanisms, such as CME.

### RET localization with cell-surface clathrin and internalization are reduced in TMEM127-depleted cells

Previous studies have shown that, in response to ligand, RET is recruited to clathrin clusters assembling on the plasma membrane and internalized into endosomal compartments via clathrin coated pits (CCP) (Crupi et al., 2020; Crupi et al., 2015; Richardson et al., 2006). To further explore the surface accumulation of RET and impairment of RET internalization in TMEM127-depleted cells, we assessed RET colocalization with cell-surface clathrin by Total Internal Reflection Fluorescence Microscopy (TIRFM), as previously described (Aguet et al., 2013; Crupi et al., 2020; Crupi et al., 2015; Hyndman et al., 2017). In Mock-KO cells, GDNF treatment significantly increased colocalization of RET and plasma membrane clathrin structures (Figure 3 A, B) and caused a significant loss of cell surface RET intensity (Figure 3A, B, Figure 4A), suggesting activated RET is being recruited to CCPs and then moving out of the TIRF field as it is internalized into the cell. In contrast, when normalized for the increased total RET intensity on the cell surface, RET-clathrin colocalization in untreated TMEM127-KO cells was significantly less than in untreated Mock-KO cells and was not increased by GDNF treatment (Figure 3A, B), suggesting impaired recruitment of RET to cell-surface clathrin irrespective of ligand stimulation. The overall intensity of cell surface clathrin puncta was not affected by GDNF treatment but was significantly less in TMEM127-KO cells compared to Mock-KO (Figure 3C). A decrease of clathrin fluorescence intensity in plasma membrane clathrin structures could suggest a decrease in the number of clathrin subunits incorporated per CCP, and thus reflect a change in clathrin assembly or size. Alternatively, as TIRF intensity decays exponentially with distance from the coverslip, a reduction of clathrin intensity in the TIRF image could also indicate an increase in membrane curvature. To distinguish between these possibilities, we used concomitant widefield epifluorescence microscopy to assess clathrin fluorescence intensity at puncta detected within the TIRF images above (Figure 3D), as this form of microscopy is not sensitive to changes in fluorescence intensity by the extent of CCP curvature generation. Clathrin fluorescence intensity in epifluorescence images was robustly reduced in clathrin puncta from TMEM127-KO cells, indicating less clathrin and smaller clusters, not more deeply invaginated structures.

**Figure 3.**
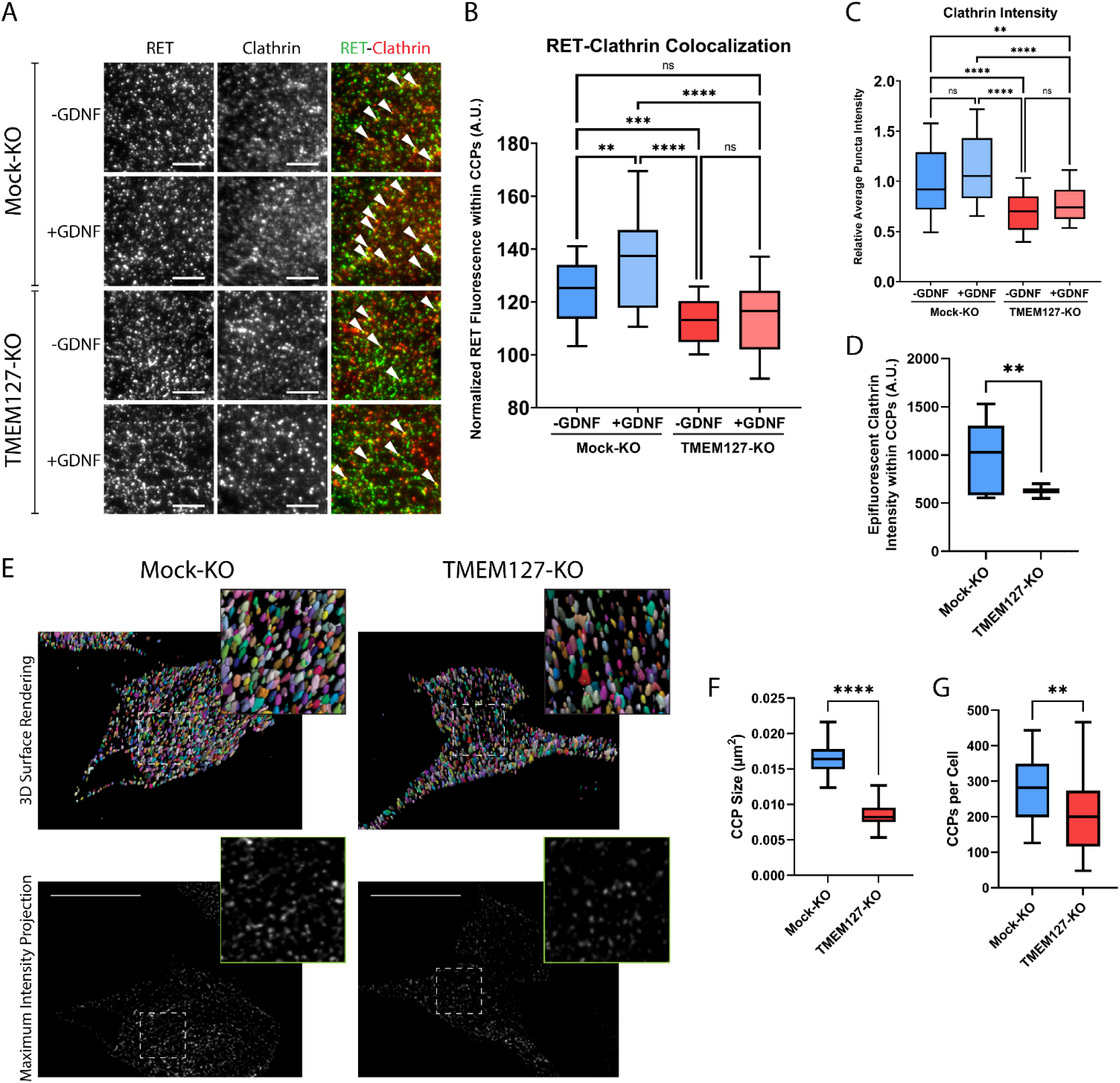
RET colocalization with cell surface clathrin is reduced and clathrin coated pits are smaller in TMEM127-depleted cells. (**A**) Representative images of RET and clathrin localization in Mock-KO and TMEM127-KO SH-SY5Y cells treated with (+) or without (-) GDNF (100 ng/ml) for 5 mins, fixed and labelled with indicated antibodies, and imaged by TIRFM. Colocalization of RET and clathrin was detected as yellow puncta (white triangles). Scale bars = 3 μm. (**B-D**) Images represented in **A** were subjected to automated detection of clathrin coated pits (CCPs) followed by quantification of the mean fluorescence intensity of RET, clathrin, or epifluorescent clathrin therein. (**B**) Mean RET fluorescence within CCPs, normalized to average RET puncta intensity, representing RET and clathrin colocalization. Data from n=30 cells representing a minimum of 5100 CCPs per condition are shown from four independent experiments. (Brown-Forsythe and Welch ANOVA tests and Dunnett’s T3 multiple comparisons test; **p<0.01, ***p<0.001, ****p<0.0001) (**C**) Average clathrin puncta intensity is shown relative to the Mock-KO (-GDNF) condition. Data from n=50 cells representing a minimum of 8700 CCPs per condition are shown from four independent experiments. (Kruskal-Wallis test and Dunn’s multiple comparisons test; **p<0.01, ****p<0.0001). (**D**) Mean epifluorescent clathrin intensity within CCPs detected by TIRF, representing intensity extending into the cell, indicating CCP depth. Data from n=34-52 cells per condition are shown from four independent experiments, representing a minimum of 8400 CCPs per condition. (Two-tailed Mann Whitney test, **p<0.01). (**E-G**) Structured illumination microscopy (SIM) imaging of EGFP-tagged clathrin light chain (EGFP-CLC) stably expressed in fixed Mock-KO and TMEM127-KO SH-SY5Y cells. (**E**) Representative images of 3D renders (upper panel) of entire cells of maximum intensity projections (lower panel) of CCPs in and near the membrane closest to the coverslip. Scale bars = 10 μm. (**F**) CCP area was measured and plotted (μm^2^) and (**G**) number of CCPs per cell was also calculated for n=26-55 cells representing a minimum of 6800 CCPs per condition from three independent experiments. (Two-tailed Mann Whitney test; **p<0.01, ****p<0.0001).

**Figure 4.**
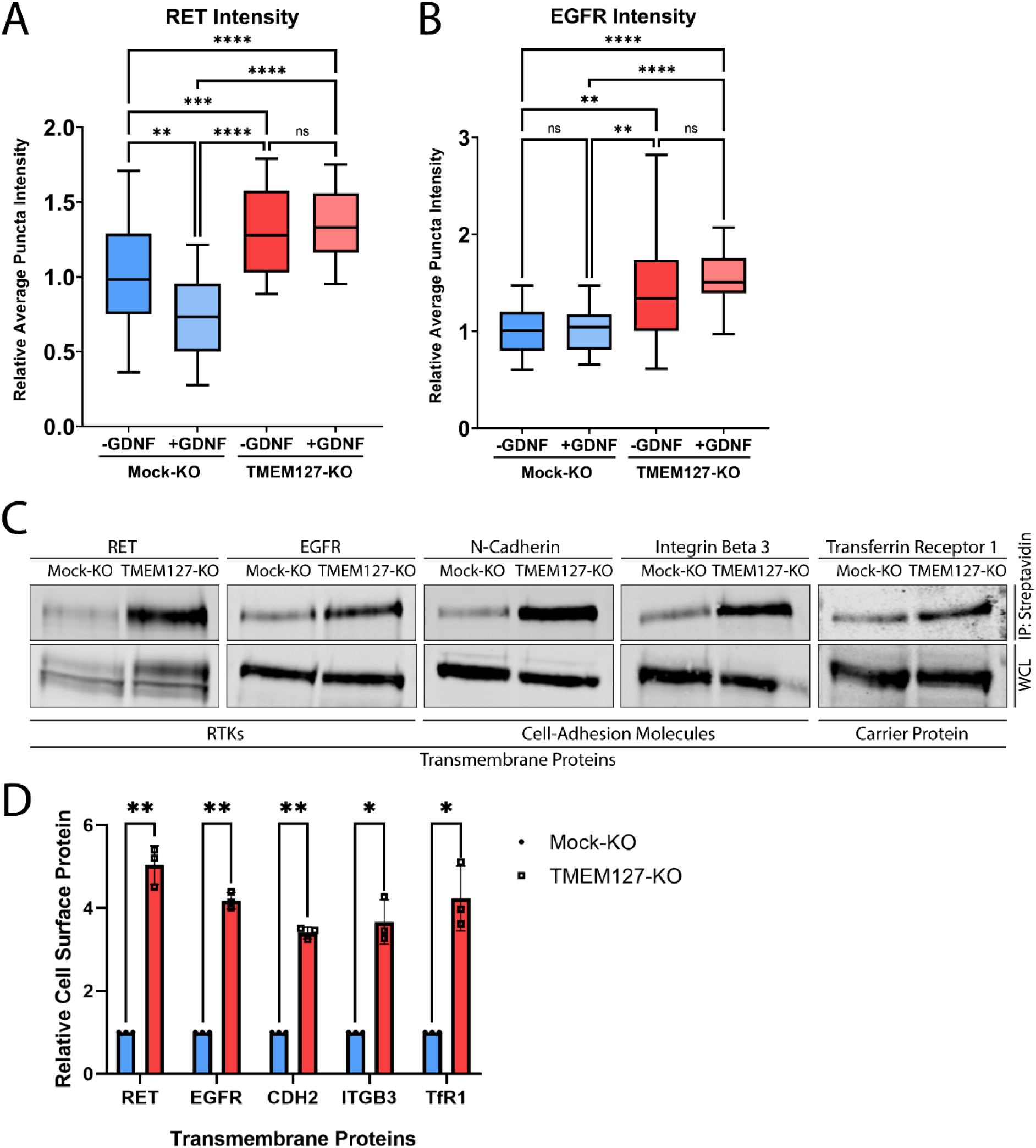
TMEM127-depletion leads to cell surface accumulation of transmembrane proteins. (**A**) TIRFM images represented in Figure 3A were subjected to automated detection of RET puncta. Average RET puncta intensity is shown relative to the Mock-KO (-GDNF) condition. Data from n=50 cells representing a minimum of 12000 RET puncta per condition are shown from four independent experiments. (Kruskal-Wallis test and Dunn’s multiple comparisons test; **p<0.01, ***p<0.001, ****p<0.0001) (**B**) TIRFM images of Mock-KO and TMEM127-KO SH-SY5Y cells were subjected to automated detection of EGFR puncta. Average EGFR puncta intensity is shown relative to the Mock-KO (-GDNF) condition. Data from n=30 cells representing a minimum of 4600 EGFR puncta per condition are shown from two independent experiments. (Brown-Forsythe and Welch ANOVA tests and Dunnett’s T3 multiple comparisons test; **p<0.01, ****p<0.0001). (**C**) Immunoblot showing the indicated total and cell surface biotinylated transmembrane proteins in Mock-KO and TMEM127-KO SH-SY5Y cells. Biotinylated proteins include RET, EGFR, N-Cadherin (CDH2), Integrin Beta-3 (ITGB3), and Transferrin Receptor-1 (TfR1). Biotinylated proteins were collected, separated, and immunoblotted as in Figure 2A. (**D**) Quantification of relative cell surface protein levels detected in C. Biotinylated cell surface protein levels (IP) were normalized to corresponding total protein (WCL) and expressed relative to Mock-KO for each protein. Three independent experiments (n=3) are shown as mean ±SD. (Two-tailed unpaired t-tests with Welch’s correction; *p<0.05, **p<0.01).

We further assessed numbers and sizes of CCPs in TMEM127 KO cells by imaging EGFP-tagged clathrin with structured illumination microscopy (SIM). Because over 60% of CCPs have been measured to be less than 200 µm in size (Wu et al., 2001), the approximate diffraction limit of a light microscope, super-resolution techniques are needed to fully resolve these structures for accurate quantitation. Therefore, we used a technique called dual iterative SIM (diSIM, SIM^2^; (Löschberger et al., 2021)) that combines a lattice illumination pattern with deconvolution and standard SIM processing to resolve lateral structures as small as 60-100 nm in the ∼525 nm emission range. Initial visual inspection of diSIM images of TMEM127-KO and Mock-KO cells expressing EGFP-tagged clathrin light chain (EGFP-CLC) suggested TMEM127-KO cells had fewer and smaller CCPs (Figure 3E). We further quantified the 2D area of over 6500 CCPs in and near the plasma membranes closest to the coverslip in TMEM127-KO and Mock-KO cells. Here, we found TMEM127-KO cells had a 2D CCP area approximately half of Mock-KO cells (0.008 µm^2^ vs. 0.016 µm^2^; Figure 3F). An area of 0.008 µm^2^ corresponds to a diameter of approximately 90 nm, within the resolving power of the diSIM technique (Löschberger et al., 2021). Finally, we also quantitated the number of CCPs per cell on a single plane near the lower membrane of TMEM127-KO and Mock-KO cells. The added resolving power of diSIM gave us confidence in reaching a more accurate count (Figure 3G). Here we found a significant decrease in CCPs in TMEM127-KO cells relative to Mock-KO cells (213 vs. 289). diSIM provides a clear indication that knockout of TMEM127 alters the size and quantity of CCPs in a cell.

Together, our data indicate that TMEM127 depletion led to smaller clathrin clusters or CCPs, suggesting that TMEM127 regulates CCP formation, assembly, and/or turnover. In contrast, we observed a significant 1.3-fold increase in intensity of cell surface RET puncta in TMEM127-KO cells relative to Mock-KO cells (Figure 4A), consistent with the cell surface accumulation of RET, but did not observe a decrease in RET puncta in response to GDNF stimulation suggesting impaired internalization. As fewer CCPs are present in TMEM127-KO cells, some of these intense RET puncta are likely not in CCPs but may represent other organization of the receptor at the plasma membrane.

Interestingly, we saw a similar increase in surface EGFR puncta intensity in TMEM127-KO cells, with a 1.4-fold increase over Mock-KO cells (Figure 4B), demonstrating that protein accumulation at the cell surface is not exclusive to RET in these cells. We extended this observation using surface biotinylation assays and showed that multiple classes of transmembrane proteins including RTKs (RET, EGFR), cell adhesion molecules (N-Cadherin, Integrin Beta-3) and carrier proteins (Transferrin Receptor-1) accumulate on the cell membrane in TMEM127-KO compared to Mock-KO cells (Figure 4C, D), suggesting a global effect caused by impaired surface protein internalization. Together, these data suggest that loss of TMEM127 impairs normal membrane processes including formation of clathrin clusters or CCPs and recruitment of cargo, resulting in altered cell membrane composition and accumulation of multiple cell surface proteins.

### TMEM127-depletion impairs assembly and maturation of clathrin coated pits

To explore perturbations of the cell membrane in TMEM127-KO cells, we used live-cell timelapse TIRF imaging of cells stably expressing EGFP-CLC (Figure 5A, Animation 1). Consistent with our fixed cell images, Mock-KO cells appeared to have more intense cell surface clathrin puncta, and these moved actively into and out of the TIRF-field at the membrane consistent with rapid formation and maturation of CCPs (Animation 1A). TMEM127-KO cells had less intense membrane clathrin puncta, which appeared less able to mature as CCPs and internalize into the cell, moving out of the TIRF field (Animation 1B).

**Figure 5.**
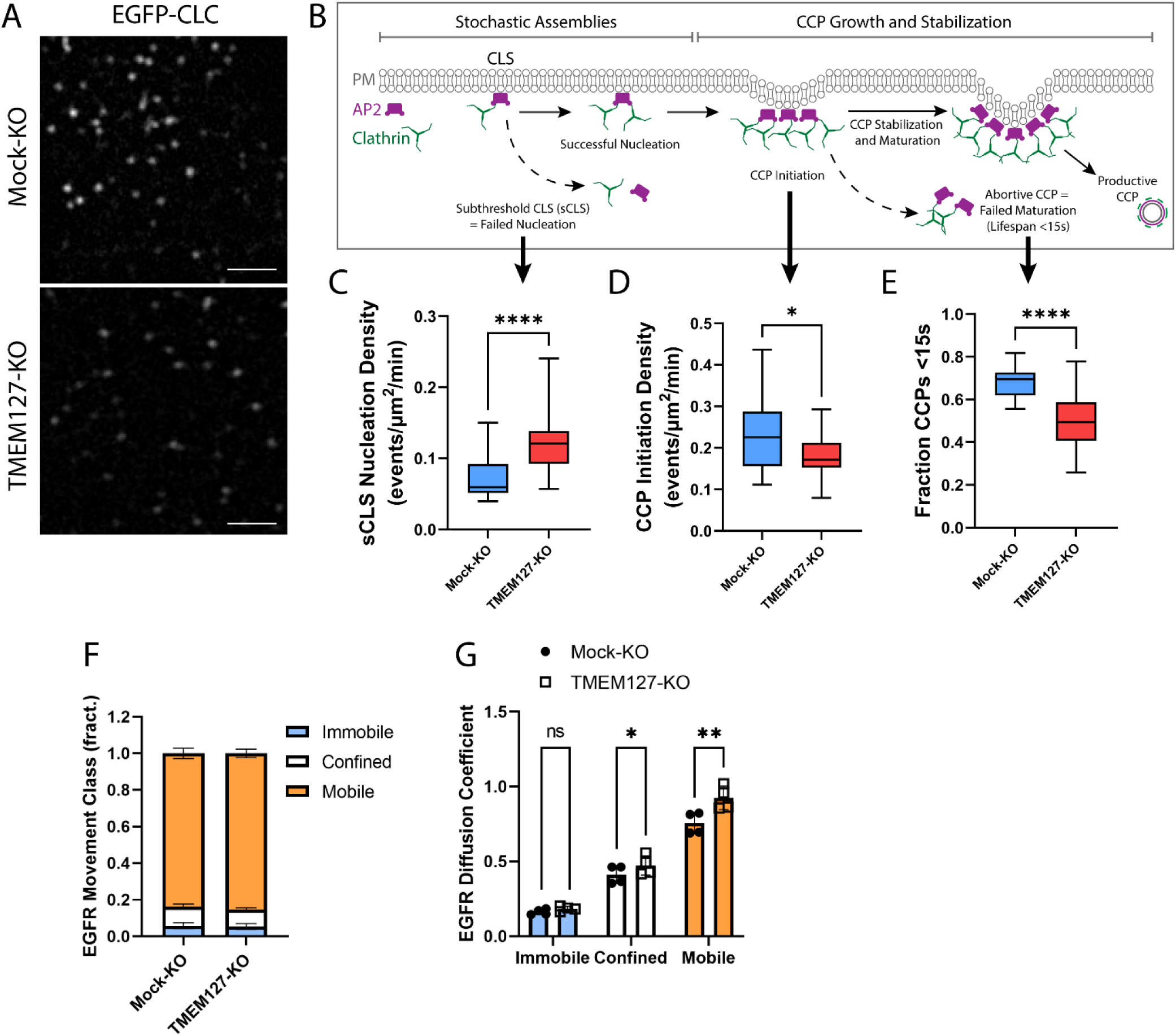
TMEM127 loss alters dynamics of CCP assembly and EGFR diffusion. Live cell TIRFM imaging of cell surface EGFP-tagged clathrin light chain (EGFP-CLC) stably expressed in Mock-KO and TMEM127-KO SH-SY5Y cells. (**A**) Representative TIRFM images of cell surface clathrin distribution. Scale bars = 2 μm. (**B**) Diagram of the stepwise process of clathrin coated pit (CCP) assembly. Plasma Membrane (PM) (grey), AP2 (purple), and clathrin (green) are indicated. Stochastic assemblies of clathrin adaptor AP2 and clathrin form clathrin labelled structures (CLS). Small structures with insufficient clathrin are unstable and may be unable to nucleate clathrin polymerization (subthreshold CLS: sCLS) and dissipate or may successfully stimulate clathrin polymerization, leading to CCP initiation. As polymerization continues, the CCP undergoes maturation and stabilization processes required to develop a productive CCP, which functions to internalize protein cargo into the cell. Unsuccessful CCP maturation leads to abortive CCPs with lifespans <15s. Steps relevant to graphs C-E are indicated with black arrows. (**C-E**) Timelapse videos of cells (5 min @ 1 frame-per-second) were subjected to automated detection of cell membrane clathrin structures, which included sCLS and bona fide CCPs. (**C**) sCLS Nucleation Density, (**D**) CCP Initiation Density, and (**E**) Fraction CCPs <15s are shown. (**C-E**) The means of multiple timelapse videos are shown. In three independent experiments, the number of total sCLS trajectories, CCP trajectories and timelapse videos (respectively) for each condition are; Mock-KO, 11040, 7988, 20, and TMEM127-KO, 15598, 8393, 28. (C-D: Two-tailed Mann Whitney test, E: Two-tailed unpaired t-test with Welch’s correction; *p<0.05, ****p<0.0001). (**F**) EGFR Movement Class shown as the mean ±SD fraction (fract.) of all EGFR tracks, as labelled by Fab-Cy3B, that exhibit mobile (orange), confined (white), or immobile (blue) behaviour. (**G**) EGFR diffusion coefficient shown is mean ±SD of immobile, confined, and mobile EGFR tracks. (**F-G**) Data from four independent experiments representing detection and tracking of minimum 500 EGFR objects. (Two-tailed paired t-tests; *p<0.05, **p<0.01)

CCPs undergo a multistage initiation, stabilization, and maturation process (Mettlen et al., 2018) (Figure 5B). Clathrin assemblies must reach a minimum threshold for nucleation of CCPs to initiate. These assemblies can result in either small short-lived (<15 s) abortive CCPs that spontaneously disassemble without cargo uptake, or longer lived productive CCPs that effectively recruit cargo for internalization and delivery to the endolysosomal system (Kaksonen and Roux, 2018; Mettlen et al., 2018). We examined CCP dynamics over time in live-cells using TIRFM and automated detection to identify diffraction-limited EGFP-CLC objects, and assessed the rates of assembly, initiation, and maturation of CCPs in Mock-KO and TMEM127-KO cells. Here, we show that TMEM127-KO cells have significantly increased stochastic assemblies of transient small subthreshold clathrin-labelled structures (sCLS) that are unable to recruit sufficient clathrin to form bona fide CCPs (Figure 5B, C). As a result of this increase in transient structures, we show a resultant decrease in the initiation of bona fide CCPs in TMEM127-KO cells compared to Mock-KO cells (Figure 5B, D).

Further, as indicated by the reduced clathrin puncta intensity at the membrane and reduced area of CCPs detected by dSIM (Figure 3C, F), the CCPs that do form in TMEM127-KO cells are smaller, potentially limiting their ability to recruit cargo, such as RET, leading the cargo proteins to accumulate on the cell surface (Figure 4 A-D). Interestingly, although there are relatively fewer CCPs formed in TMEM127-KO cells (Figure 5D), a smaller fraction of these had lifespans less than 15 s, suggesting CCPs that are able to overcome early assembly defects due to TMEM127 loss are able to mature to productive CCPs more efficiently (Figure 5E). The overall increase in sCLS and decrease in CCPs, suggests that TMEM127 depletion may reduce the potential for clathrin to form or stabilize CCPs capable of internalization and compromise the maturation of productive CCPs, thereby leading to impaired internalization of RET and other cell surface proteins.

The defect in cargo recruitment to CCPs (Figure 3A, B) and CCP assembly (Figure 3C-G, 5A-E) in TMEM127-KO cells could reflect changes specific to cargo engagement within CCPs or broader changes in membrane dynamics, such as protein and lipid mobility. To determine if loss of TMEM127 resulted in broader impacts on membrane dynamics, we examined the mobility of EGFR by single-particle tracking, using strategies we recently developed and characterized (Sugiyama et al., 2023).

Importantly, labeling of EGFR with a Fab antibody fragment for single particle tracking in the absence of ligand stimulation allows study of EGFR mobility that is largely clathrin independent (Sugiyama et al., 2023). TMEM127-KO cells exhibited a similar distribution of the population of EGFR in immobile, confined, and mobile cohorts (Figure 5F), suggesting that TMEM127 does not disrupt capture of a minor subset of EGFR within tetraspanin nanodomains that occurs in the absence of ligand (Sugiyama et al., 2023). However, consistent with broader disruption of membrane dynamics upon loss of TMEM127, we observed an increase in the diffusion coefficient of the confined and mobile fractions, in particular the latter (Figure 5G). This predicts that loss of TMEM127 may broadly impact membrane dynamics, suggesting that defects in receptor recruitment to CCPs and perturbations of CCP assembly that may lead to accumulation of membrane proteins at the cell surface in TMEM127-KO cells may result in part from broad alterations in membrane dynamics and/or receptor mobility that impact the formation of CCPs and their recruitment of diverse cargo.

### Clathrin adaptor recruitment is not TMEM127 dependent

Recruitment of RET to CCPs requires interaction with the clathrin adaptor AP2 (Crupi et al., 2015). We used proximity ligation assays (PLA) to identify and quantify direct protein interactions *in situ* to assess association of endogenous AP2 with RET and clathrin in SH-SY5Y cells. In untreated Mock-KO cells, RET-AP2 interaction was minimal but was significantly increased by GDNF treatment, as indicated by increased PLA puncta (Figure 6A), suggesting RET recruitment as cargo to CCPs. Association of RET and AP2 was significantly greater in TMEM127-KO cells, consistent with the higher RET protein levels at the cell membrane, and was further increased by GDNF treatment, suggesting that the RET-AP2 association was not TMEM127 dependent. We saw robust association of clathrin and AP2 in untreated Mock-KO cells (Figure 6B). GDNF treatment significantly reduced association, consistent with maturation of productive CCPs and subsequent uncoating of clathrin coated vesicles and delivery of cargo to endosomes. In comparison, TMEM127-KO cells had significantly greater constitutive association of AP2 and clathrin and this was not affected by GDNF. We did not detect association of AP2 with TMEM127 under any condition using PLA (not shown), suggesting TMEM127 is not closely associated with clathrin labelled structures. Together, our data suggest that, in TMEM127-KO cells, AP2-clathrin complexes accumulate but are unable to efficiently support the formation and/or assembly of CCPs to facilitate internalization of cargo proteins. This is consistent with the observation that TMEM127-KO cells have significantly higher sCLS nucleation rates, while exhibiting decreased formation of bona fide CCPs (Figure 5C, D). It follows that the increased proximity of RET and AP2 in TMEM127-KO cells may reflect increased cell surface levels of RET that may engage with endocytic adaptors, but that these interactions infrequently lead to formation or stabilization of bona fide CCPs. These data suggest that TMEM127 depletion does not directly affect ability of adaptors to associate with clathrin or cargo but may reflect an integral membrane defect that alters normal membrane protein movement and abilities to assemble or stabilize larger membrane protein complexes.

**Figure 6.**
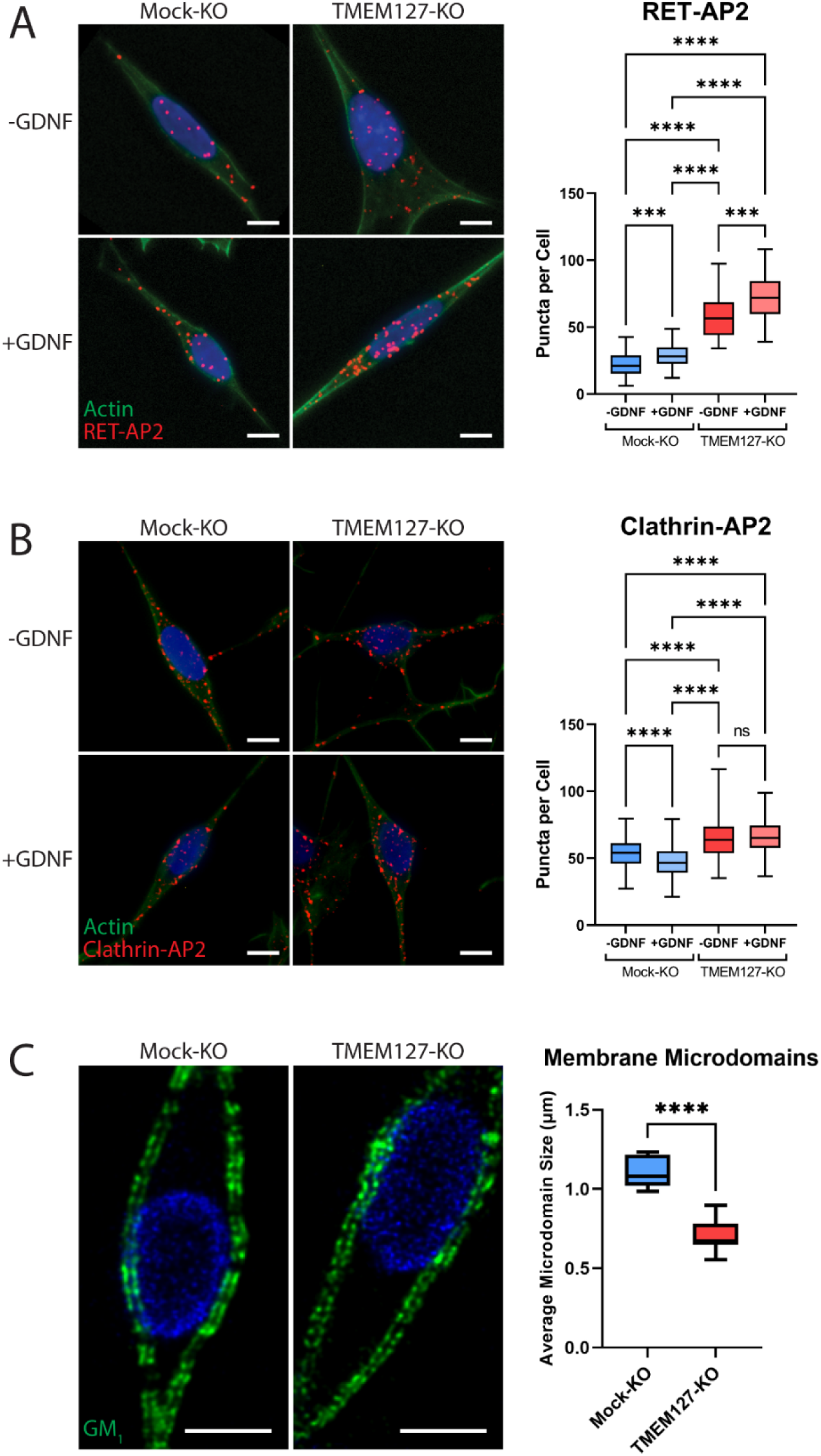
Enhanced recruitment of endocytic adaptors and disrupted membrane microdomains in TMEM127-depleted cells. (**A** and **B**) Proximity ligation assay (PLA) of Mock-KO and TMEM127-KO SH-SY5Y cells treated with (+) or without (-) GDNF (100 ng/ml) for 5 mins. Representative immunofluorescence images (left) of RET-AP2 (**A**) and Clathrin-AP2 (**B**) interactions (red puncta). Cells were stained with Phalloidin actin filament stain (green) and Hoechst nuclear stain (blue). Quantification of red puncta per cell (right), indicating individual protein interactions in each condition, are shown. Data from a minimum of 100 cells (-GDNF conditions) (**A**), 300 cells (+GDNF conditions) (**A**), or 500 cells (all conditions) (**B**) are shown. (Kruskal-Wallis test and Dunn’s multiple comparisons test; ***p<0.001, ****p<0.0001). Scale bars = 5 μm. (**C**) Immunofluorescence confocal images of Mock-KO and TMEM127-KO SH-SY5Y cells (left) stained with cholera toxin subunit B, which binds ganglioside GM_1_-positive lipid microdomains (membrane rafts) (green), and Hoechst nuclear stain (blue). Quantification of average lipid raft size (right) of n=11-12 cells per condition are shown, representing a minimum of 500 lipid rafts per condition. (Two-tailed unpaired t-test; ****p<0.0001). Scale bars = 5 μm.

### TMEM127 depletion alters membrane microdomain organization

Given the results of the EGFR single-particle tracking experiments that suggest broader disruption of membrane dynamics upon loss of TMEM127 (Figure 5F, G), we investigated whether loss of TMEM127 could more broadly impact membrane organization by assessing distribution of lipid-rich membrane microdomains (membrane rafts). In Mock-KO cells stained for ganglioside G_M1_, which selectively partitions into membrane rafts (Sonnino et al., 2007), stained lipid microdomains were large and continuous (Figure 6C). In TMEM127-KO cells, staining was fragmented and microdomains were significantly smaller, suggesting inability to assemble large lipid clusters and raft-associated proteins in the surface membrane. These data suggest that TMEM127-depletion alters membrane dynamics, limiting formation of multiple membrane structures.

### Decreased internalization reduces RET degradation, increasing RET half life

Internalization of activated RET into the endolysosomal system is required for downregulation of RET signaling and eventual degradation (Crupi et al., 2020; Hyndman et al., 2017). As RET internalization was reduced in TMEM127-KO cells, we assessed whether this might impair RET degradation by examining RET protein expression levels over time following GDNF treatment in cells treated with cycloheximide (CHX) to inhibit new protein synthesis. In GDNF-treated Mock-KO cells, RET was rapidly degraded, with a half-life under 2 hours and less than 9% RET protein remaining after 12 hours (Figure 7). In these cells, TMEM127 was shown to have a half-life of approximately 5 hours. In contrast, RET degradation was significantly slower in TMEM127-KO cells, with a half-life of approximately 6 hours and over 43% of RET remaining after 12 hours (Figure 7). These data suggest that impaired RET internalization due to altered membrane dynamics significantly reduces RET degradation, contributing to RET accumulation on the cell surface.

**Figure 7.**
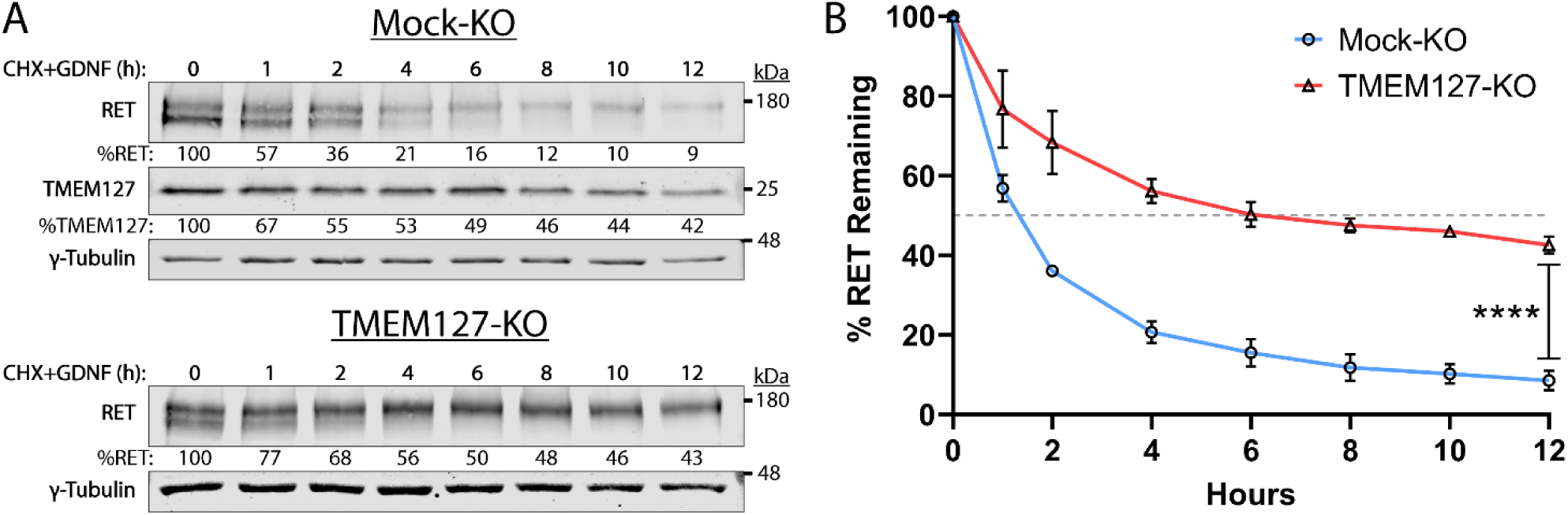
RET half-life is increased when TMEM127 is depleted. (**A**) Mock-KO (upper panels) and TMEM127-KO (lower panels) SH-SY5Y cells were treated with cycloheximide (CHX) and GDNF (100 ng/ml) for indicated times to evaluate RET and TMEM127 protein levels remaining over time. Whole cell lysates were separated by SDS-PAGE and immunoblotted for the indicated proteins. γ-Tubulin was used as a loading control. RET and TMEM127 protein remaining at each time point, relative to untreated levels (0 h), is indicated below each lane. (**B**) Quantification of RET protein remaining over time as shown in **A**. The dashed line indicates 50% RET remaining. The means ±SD at each timepoint of three independent experiments (n=3) are shown. (Two-way ANOVA with Šídák’s multiple comparisons test; ****p<0.0001).

### RET accumulation promotes increased downstream signalling

Overexpression or cell surface accumulation can frequently cause constitutive activation of growth factor receptors, conferring ligand insensitivity for modulating the intensity and duration of their downstream signals (Du and Lovly, 2018). We assessed RET activation and signaling over time in Mock-KO and TMEM127-KO cells in response to GDNF. In Mock-KO cells, RET and its downstream signaling intermediates AKT, ERK and the mTOR substrate S6 were minimally phosphorylated in the absence of GDNF and showed significantly increased phosphorylation over time in response to GDNF treatment (Figure 8A, B), consistent with our previous studies (Crupi et al., 2020; Richardson et al., 2006). In contrast, RET, AKT, ERK and S6 were constitutively phosphorylated in TMEM127-KO cells. GDNF enhanced RET phosphorylation after 5 min but, unlike Mock-KO cells, the effect was not further increased by prolonged treatment times. Phosphorylation of AKT, ERK and S6 was not significantly increased by GDNF treatment in TMEM127-KO cells (Figure 8A, B). Further, we showed that the constitutive phosphorylation of RET, AKT, ERK, and S6 in TMEM127-KO cells could be blocked by RET inhibition using the multikinase inhibitor Vandetanib, and selective RET inhibitors Selpercatinib, and Pralsetinib (Figure 8C), which are clinically approved for treatment of RET-associated cancers (Mulligan, 2019). GDNF-mediated phosphorylation in Mock-KO cells was also blocked by RET inhibition (Figure 8-figure supplement 1). Together, our data show that cell surface accumulation of RET can promote constitutive phosphorylation and activation of multiple downstream signaling pathways.

**Figure 8.**
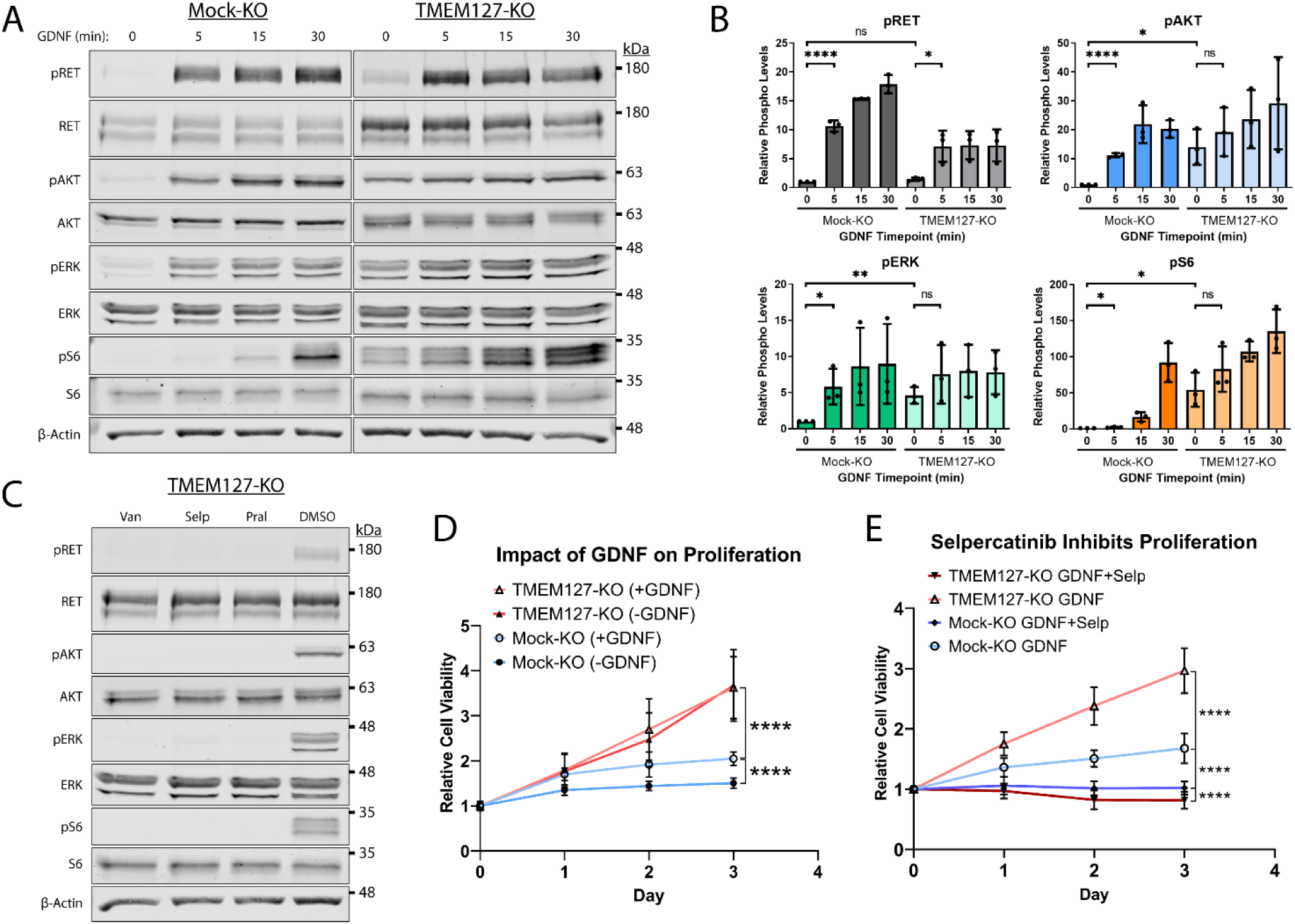
Constitutive RET-mediated signalling drives proliferation and confers sensitivity to Selpercatinib. (**A**) Mock-KO and TMEM127-KO SH-SY5Y cells were serum starved and treated with GDNF for the indicated times. Whole cell lysates were separated by SDS-PAGE and immunoblotted for the indicated Phospho and Total proteins. β-Actin was used as a loading control. (**B**) Quantification of phosphorylated protein levels as shown in **A** relative to Mock-KO 0 h condition for each protein. The means ±SD of three independent experiments (n=3) are shown (Two-tailed unpaired t-tests; *p<0.05, **p<0.01, ****p<0.0001). (**C**) Expression of the indicated Phospho and Total proteins in TMEM127-KO SH-SY5Y cells treated with DMSO, 5 μM Vandetanib (Van), Selpercatinib (Selp), or Pralsetinib (Pral) (1 h), and immunoblotted as in **A**. Constitutive phosphorylation of indicated proteins was abrogated by all three inhibitors. (**D**) Cell viability of Mock-KO and TMEM127-KO SH-SY5Y cells over time in the presence or absence of GDNF, relative to Day 0 was measured by MTT assay as an indicator of cell proliferation. (**E**) Cell viability of Mock-KO and TMEM127-KO SH-SY5Y cells in the presence of GDNF (100 ng/ml) alone or GDNF+Selpercatinib (0.1 μM), relative to Day 0, as above. (**D and E**) Three independent experiments, for each graph, with 18 replicates per condition and timepoint (n=54) are shown as mean ±SD (Two-way ANOVA with Tukey’s multiple comparisons test; ****p<0.0001).

### RET drives proliferation in TMEM127-depleted cells

Oncogenic activating RET mutations are drivers of proliferation and tumour growth in PCC (Mulligan, 2019; Neumann et al., 2019; Toledo et al., 2017). Here, we assessed whether RET constitutive activation caused by cell surface accumulation in response to TMEM127 loss could also promote these processes. We showed that GDNF treatment significantly increased proliferation of Mock-KO cells over a 3-day period (Figure 8D). TMEM127-KO cells proliferated significantly faster than Mock-KO cells, irrespective of GDNF treatment (Figure 8D), consistent with increased RET accumulation leading to ligand independent RET activity and proliferation. To confirm that proliferation was RET-mediated and explore the impact of RET inhibition in TMEM127-KO cells, we assessed proliferation in the presence and absence of the selective RET inhibitor Selpercatinib at a concentration sufficient to block RET phosphorylation (0.1 μM, Figure 8-figure supplement 2).

Selpercatinib significantly reduced GDNF-dependent proliferation of Mock-KO cells. In TMEM127-KO cells, RET inhibition abrogated the increase in proliferation, seen above, and further significantly reduced viability (∼20%) compared to Mock-KO cells (Figure 8E). Vehicle alone (DMSO) did not affect proliferation under any conditions (Figure 8-figure supplement 3). Our data demonstrate that proliferation of TMEM127-KO cells was RET-dependent and suggest a previously unrecognized sensitivity of TMEM127-depleted cells to RET inhibition.

## Discussion

The RET receptor and the TMEM127 integral membrane protein are established PCC susceptibility genes (Dahia, 2014; Mulligan, 2019; Neumann et al., 2019). RET oncogenic mutations give rise to PCC in patients with MEN2 and are found in up to 5% of sporadic PCC, where they promote constitutive RET activation and downstream signaling (Horton et al., 2022; Le Hir et al., 2000; Mulligan, 2019; Takaya et al., 1996a; Takaya et al., 1996b). In contrast, TMEM127 acts as a tumour suppressor, and loss of function mutations have been linked primarily to increased mTOR signaling (Deng et al., 2018). Although RET and WT-TMEM127 localize to similar endosomal compartments, we show that colocalization is lost in the presence of PCC-associated mutants that disrupt TMEM127 membrane insertion (Qin et al., 2010). Endosomes are reduced in size and number in the absence of WT-TMEM127, suggesting defects in the processes required for vesicle formation and internalization and trafficking of proteins from the plasma membrane (Figure 9).

**Figure 9.**
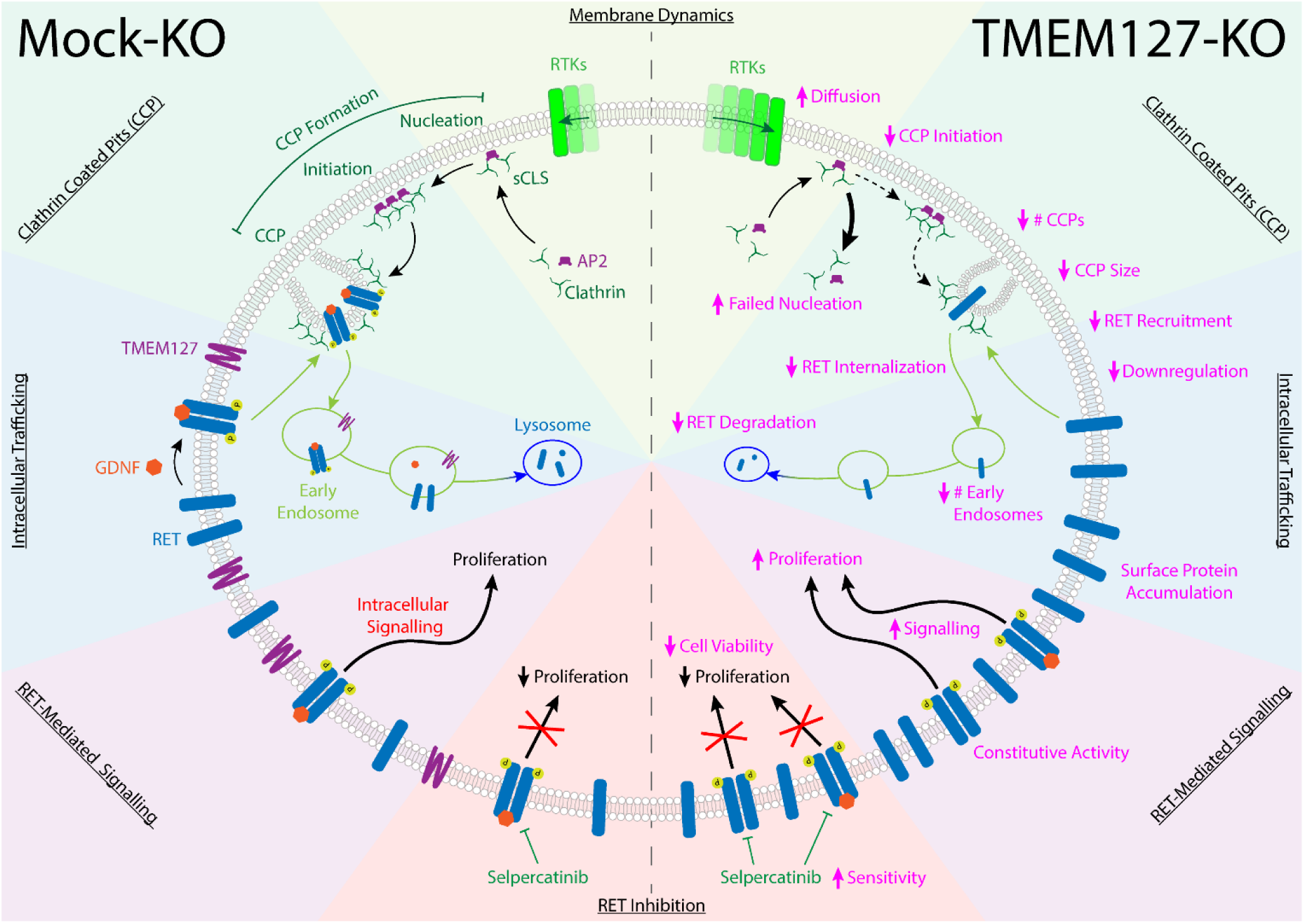
TMEM127 loss drives transformation through altered RET location and activity in PCC. TMEM127 depletion alters membrane diffusion and impairs formation of productive clathrin coated pits (CCP) for internalization of membrane receptors. CCPs are fewer, smaller, and less able to recruit cargo, resulting in accumulation of transmembrane proteins, including RET on the cell surface. As a result, RET downstream signaling is enhanced and degradation is reduced leading to increased cell proliferation. The selective RET inhibitor Selpercatinib abrogates RET-mediated signals and blocks cell proliferation.

Using a CRISPR TMEM127-KO in the neuroblastoma cell line SH-SY5Y, we noted significantly increased levels of endogenously expressed RET protein, which appeared primarily as the higher molecular weight fully glycosylated form found at the cell surface (Richardson et al., 2006; Richardson et al., 2012; Takahashi et al., 1991). Using surface biotin labelling and TIRFM in fixed and live cells we showed that RET internalization was impaired, and that WT RET accumulated on the cell surface in the absence of TMEM127, escaping degradation (Figure 9). Notably, accumulation of surface receptors was not unique to RET as we also saw increased intensity of EGFR puncta and increased membrane levels of other receptors and adhesion molecules in the absence of TMEM127 in both SH-SY5Y and HEK293 cells, consistent with a global defect of membrane dynamics that could also block internalization of membrane proteins. As a result of increased density in the plasma membrane in SH-SY5Y cells, RET is constitutively active, promoting stimulation of multiple downstream signaling pathways. Importantly, treatment with selective RET inhibitors abrogated these signals including mTOR signaling, suggesting that, in this model, the increase in mTOR signaling seen in response to TMEM127 loss is a direct response to RET activity.

As both RET and TMEM127 have been previously shown to internalize through clathrin mediated endocytosis (Crupi et al., 2020; Crupi et al., 2015; Flores et al., 2020; Richardson et al., 2006) we assessed the impact of TMEM127 depletion on membrane clathrin distribution and its effects on cargo internalization. Despite increased RET puncta intensity on the cell surface, RET localization to clathrin puncta was reduced and was not responsive to GDNF, consistent with impairment of clathrin mediated internalization of RET (Figure 9). The formation of CCPs is a multistep process involving recruitment of clathrin adaptors and stepwise assembly of clathrin labelled structures at the cell membrane for recruitment of cargo proteins and assembly of productive CCPs (Mettlen et al., 2018).

Transient small clathrin labelled structures (sCLSs) are unstable and may not recruit sufficient clathrin or cargo to achieve a threshold size and progress further to initiate bona fide CCPs (Mettlen et al., 2018). Once sufficient clathrin is recruited for initiation, CCPs may mature to pinch off clathrin coated vesicles from the membrane, but a subset fail to mature and are aborted (Kadlecova et al., 2017; Mettlen et al., 2018). Our single particle tracking studies suggest that mobility of membrane cargo proteins like EGFR, is enhanced in TMEM127 KO cells, which may decrease capture of cargo at stochastic clathrin assemblies and limit their progression to successful nucleation and initiation of bona fide CCPs. Consistent with this, our data indicate that TMEM127-depletion increased the frequency of the sCLSs but significantly reduced their ability to reach the clathrin threshold size and initiate bona fide CCPs. CCPs that did form were smaller, contained less clathrin, and were able to recruit less RET cargo than in Mock-KO cells. TMEM127 loss did not impair RET-AP2 or AP2-clathrin association, suggesting that key protein interactions required for internalization were largely intact (von Zastrow and Sorkin, 2021). However, we saw decreased capture of RET within bona fide CCPs, formation of CCPs was reduced, and those that formed smaller in TMEM127 deficient cells, suggesting that altered membrane dynamics, increasing the diffusability of membrane proteins and impairing capture of sufficient cargo-adaptor complexes into nascent CCPs, could be limiting the recruitment of sufficient cargo and coalescence of CCPs, consistent with previous studies that link membrane tension and CCP formation (Djakbarova et al., 2021; Saleem et al., 2015). Together, our data suggest that accumulation of RET and other transmembrane proteins at the cell surface in the absence of functional TMEM127 is due to reduced efficiency of clathrin recruitment to CCP, limiting capacity for cargo internalization and reducing CME, and that this correlates with the global disruption of membrane protein composition, protein complex assembly, and organization in other domains, such as membrane rafts. Interestingly, our preliminary studies have suggested that one outcome of this may be impairment of the formation of other membrane-associated complexes and processes, such as recruitment of ubiquitination machinery (Guo et al., 2023), contributing to the TMEM127 phenotype and highlighting the broader implications of altered membrane partitioning.

As a result of impaired internalization in TMEM127 deficient SH-SY5Y cells, RET protein accumulates on the cell surface and degradation is reduced, leading to sustained downstream pro-proliferative signals. We showed that these pathways can be blocked using selective RET inhibitors, suggesting that targeting RET may be a valuable therapeutic strategy in the subset of PCC tumours where RET mutations have not been recognized but RET expression is detected. In support of this, in recent preliminary studies, we have shown that xenografts of TMEM127 KO SH-SY5Y cells produce larger tumors in nude mice, but that growth is reduced by treatment with the RET inhibitor Selpercatinib (Guo et al., 2023). Interestingly, TMEM127 mutations have also been identified as drivers in other cancers, notably renal cell carcinoma, where RET is not highly expressed. In these cancers, we would predict that TMEM127 loss has similar effects on membrane dynamics, membrane protein accumulation and CCP formation to those seen in our model and that accumulation of other growth factor receptors, such as MET, on the cell surface will lead to transformation (Marona et al., 2019; Rhoades Smith and Bilen, 2019). Notably, TMEM127 mutations or reduced expression are also seen at low frequencies in a number of other tumour types (e.g. endometrial, liver, breast, kidney, ovarian cancers (Tate et al., 2019)) where TMEM127 contributions have not yet been specifically recognized, suggesting that aberrant accumulation and activation of growth factor receptors could also impact these tumours. Together, our data suggest a novel paradigm for oncogenic transformation in

TMEM127 mutant tumours. We predict that altered membrane dynamics blocks normal internalization and degradation of key wildtype growth-promoting receptors, which then act as oncogenes to drive tumorigenesis in a cell type-specific fashion. Further, we predict that these oncogenes will provide valuable alternative therapeutic targets that have not been previously explored in TMEM127-mutant cancers.

Importantly, the changes in membrane dynamics and aberrant accumulation of membrane proteins in response to TMEM127 depletion are not exclusively oncogenic but may have other pathological implications. Consistent with our data showing RET accumulation on the cell surface, TMEM127 KO mice have increased insulin sensitivity and upregulation of insulin dependent AKT signaling (Srikantan et al., 2019), which may suggest excess surface accumulation of insulin receptors and resulting enhanced signaling in the liver in this model. Notably, recent studies have suggested that TMEM127 may have immune modulatory roles. In Salmonella-infected but not normal antigen presenting cells, TMEM127 acts in recruitment of protein complexes that promote downregulation of MHC-II and suppression of T-cell activation, while depletion of TMEM127 in infected cells increases MHC-II surface expression and reduces innate immune responses (Alix et al., 2020). Similarly, TMEM127 depletion leads to MHC-I accumulation on the cell membrane in AML cell models, promoting enhanced recruitment of CD8^+^ T cells (Chen et al., 2023). Taken together, these data suggest that the outcomes of TMEM127 depletion may be cell type as well as tumour-type and stage specific, depending on the cell membrane proteomic repertoire. Further, TMEM127 depletion may promote oncogenic signaling in tumour initiation and growth, as we observed in PCC, or potentially contribute to alterations in the tumour immune environment in progression of other cancer models.

These observations suggest that TMEM127 may have broad cellular roles in maintaining balance in the membrane proteome and suggest a generalized role for TMEM127 as a facilitator of diverse membrane protein complexes regulating localized membrane diffusion and promoting stability of complex assemblies to regulate membrane organization.

## Materials and Methods

### Cell Culture

Human Retinal Pigment Epithelial ARPE-19 cells (CRL-2302, ATCC, Manassas, VA, USA, RRID:CVCL_0145), were cultured in Ham’s F-12 medium (ThermoFisher Scientific, Waltham, MA, USA) with 10% Fetal Bovine Serum (FBS; Sigma Aldrich, Oakville, ON, CA). TMEM127-KO and Mock-KO were generated in SH-SY5Y neuroblastoma cells (CRL-2266, ATCC, RRID:CVCL_0019) and HEK293 (CRL-1573, ATCC, RRID:CVCL_0045) using CRISPR-Cas9 as previously described (Deng et al., 2018). SH-SY5Y and HEK293 cells were maintained in Dulbecco’s Modified Eagles Medium (DMEM; Sigma Aldrich) with 10% and 5% FBS respectively, 10 µg/mL Ciprofloxacin, and 2.5 µg/mL Puromycin. Cell lines were authenticated by STR profiling (Centre for Applied Genomics, SickKids, Toronto, ON). EGFP-tagged clathrin light chain (EGFP-CLC) expressing variants of these cells were generated through lentiviral transduction, as previously described (Crupi et al., 2020), and selected in 100 μg/ml Hygromycin B (BioShop, Burlington, ON). Prior to each experiment, endogenous RET expression in SH-SY5Y was stimulated with 5 μM retinoic acid (MilliporeSigma, Oakville, ON, CA) for 12-18 h. Where appropriate, RET activation was induced with 100 ng/ml GDNF (Alomone Labs, Israel) for the indicated times.

### Antibodies, Stains, and Inhibitors

Total RET (#14556, RRID:AB_2798509), pRET (#3221, RRID:AB_2179887), AKT (#9272, RRID:AB_329827), pAKT (#4058, RRID:AB_331168), S6 (#2317, RRID:AB_2238583), and pS6 (#2211, RRID:AB_331679) antibodies were from Cell Signaling Technologies (Beverly, MA, USA). Antibodies to RET51 (#sc-1290, RRID:AB_631316), β-Actin (#sc-47778, RRID:AB_626632), EGFR (#sc-120, RRID:AB_627492), ERK1 (#sc-94, RRID:AB_2140110), pERK (#sc-7383, RRID:AB_627545), EGFP (#sc-9996, RRID:AB_627695), and EEA1 (#sc-137130, RRID:AB_2246349) were from Santa Cruz Biotechnology (Dallas, TX, USA). The anti-γ-Tubulin (#T6557, RRID:AB_477584) antibody was from Sigma Aldrich. Clathrin Heavy Chain antibodies (#ab2731 (RRID:AB_303256) and #ab21679 (RRID:AB_2083165)), RET (# ab134100, RRID:AB_2920824), and Alpha-adaptin (ab2730, RRID:AB_303255) were from Abcam (Cambridge, UK). The TMEM127 (#A303-450A, RRID:AB_10952702) antibody was from Bethyl Laboratories (Fortis Life Sciences, Waltham, MA, USA). Western blot IRDye secondary antibodies were from LI-COR Biosciences (Lincoln, NE, USA). Alexa Fluor-conjugated secondary antibodies (Alexa-488, -546, -594, and -647) and Phalloidin (Alexa-488) were from ThermoFisher Scientific. Cyanine Cy3- and Cy5-conjugated AffiniPure fluorescent secondary antibodies were from Jackson ImmunoResearch Laboratories Inc. (West Grove, PA, USA). Nuclear stains used include Hoechst 33342 (ThermoFisher Scientific) and DAPI 10236276001 (Roche Diagnostics, Mannheim, Germany). Kinase inhibitors used were Vandetanib (AstraZeneca PLC, Mississauga, ON), Selpercatinib and Pralsetinib (Selleckchem, Houston, TX, USA).

### Expression Constructs and Transfection

Expression constructs for WT RET51, GFRα1 and FLAG- or EGFP-tagged TMEM127 WT and mutants (C140Y and S147del) have been previously described (Hyndman et al., 2017; Qin et al., 2010; Richardson et al., 2012). EGFP-tagged rat clathrin light chain (Ehrlich et al., 2004) was cloned into pLenti CMV and sequence verified.

Cells were transiently transfected with the indicated constructs using Lipofectamine 2000 (Invitrogen, Burlington, ON, CA), according to the manufacturer’s instructions (Crupi et al., 2020). For ARPE-19 cells, medium was changed 6 h after transfection and cells were incubated at 37°C for 48 h prior to fixation. For SH-SY5Y and HEK293 cells, medium was changed 5 h after transfection and cells were incubated at 37°C overnight (16h) prior to harvesting.

### Protein Isolation and Immunoblotting

Cells were washed with phosphate-buffered saline (PBS) and protein was harvested with lysing buffer (20 mM Tris-HCl (pH 7.8), 150 mM NaCl, 1 mM sodium orthovanadate, 1% Igepal, 2 mM EDTA, 1 mM phenylmethylsulfonyl fluoride, 10 µg/mL aprotinin, and 10 µg/mL leupeptin), as described (Crupi et al., 2020; Hyndman et al., 2017). Proteins were separated by SDS-PAGE and transferred to nitrocellulose membranes, as previously described (Crupi et al., 2020; Hyndman et al., 2017). Blots were probed with indicated primary (1:1000-1:5000 dilution) and secondary IRDye antibodies (1:20000) and visualized using the Odyssey DLx Imaging System (LI-COR Biosciences).

Immunoblot band intensity quantification was performed using ImageStudio Lite Ver 5.2 software (LI-COR Biosciences).

### Biotinylation assays

Biotinylation of cell-surface protein was performed as described (Crupi et al., 2020; Reyes-Alvarez et al., 2022). Briefly, cell surface proteins were labelled with biotin and either harvested immediately (simple cell-surface biotinylation assay) or cells were treated with GDNF for the indicated times to induce RET internalization (biotinylation internalization assay). Following internalization, remaining biotin was removed from the cell surface using MeSNa stripping buffer. Cells were lysed and biotinylated proteins immunoprecipitated with streptavidin-conjugated agarose beads (ThermoFisher Scientific) overnight at 4°C with agitation. Biotin-labelled proteins were collected by centrifugation at 2000×g, washed 4 times with lysing buffer and resuspended in Laemmli buffer prior to SDS-PAGE, as previously described (Crupi et al., 2020).

### Degradation assays

Serum-starved cells were treated with serum-free media containing GDNF and 100 μg/ml cycloheximide for the indicated times. Proteins were collected for immunoblotting, as above.

### Proliferation assays

Cell proliferation was evaluated through a 3-(4,5-Dimethylthiazol-2-yl)-2,5-diphenyltetrazolium bromide (MTT) reduction assay. Briefly, 1.5 × 10^4^ cells/well were seeded in 96-well plates and allowed to proliferate for indicated times. MTT (Sigma Aldrich) was added to a final concentration of 0.45 mg/ml for 16 h. Acidified 20% SDS was added, and cells were incubated at room temperature for 4 h with shaking. Formazan product was quantified by absorbance at 570 nm and normalized to blank wells lacking cells. Absorbance relative to the 0 h timepoint for each condition indicated relative cell-viability.

### Immunofluorescence and Microscopy

ARPE-19 cells were seeded onto poly-L-lysine coated coverslips for transfection. SH-SY5Y cells were seeded onto bovine collagen 1 (Corning, Oneonta, NY) coated coverslips, or into ibiTreat µ-slide VI 0.4 channel slides (Ibidi, Munich, Germany). Cells were fixed with 4% paraformaldehyde, permeabilized with 0.2% Triton X-100, and blocked in 3% BSA. Cells were incubated with the indicated primary antibodies (1:200-1:400) overnight at 4°C, followed by Alexa-Fluor/Cyanine secondary antibodies (1:2000) and indicated stains, Phalloidin (1:40), DAPI/ Hoechst 33342 (1:5000-1:10000). Lipid Raft staining was performed with the Vybrant® Alexa-Fluor 488 Lipid Raft Labelling Kit according to the manufacturer’s instructions (ThermoFisher Scientific). Coverslips or channel slides were then mounted with MOWIOL-DABCO mounting media (Sigma-Aldrich) and stored at 4°C.

Proximity Ligation Assay (PLA) was performed using the Duolink® PLA reagents according to the manufacturer’s instructions (Sigma Aldrich). An EVOS M7000 imaging system (ThermoFisher Scientific) with a 40x air objective was used to acquire immunofluorescent images of the PLA. Particles were analyzed using ImageJ software (US National Institutes of Health, Bethesda, MD, RRID:SCR_003070).

A Leica TCS SP8 confocal microscope system (Leica Microsystems, Wetzlar, HE, Germany) with a 63x/1.40 oil objective lens was used to acquire 0.9 μm optical sections. Images were deconvoluted and ImageJ software (US National Institutes of Health) was used to analyze colocalization and normalize channel intensities to generate the final images.

Total Internal Reflection Fluorescence Microscopy (TIRFM) was performed using a Quorum (Guelph, ON, CA) Diskovery microscope, comprised of a Leica DMi8 microscope equipped with a 63×/1.49 NA TIRF objective with a 1.8× camera relay (total magnification 108×). Images were acquired using a Zyla 4.2-megapixel sCMOS camera (Oxford Instruments) with 488-, 561-, or 637-nm laser illumination and 527/30, 630/75, or 700/75 emission filters. Fixed-cell TIRFM imaging was performed at room temperature with samples mounted in PBS. Live-cell imaging experiments were performed with cells at constant 37°C during imaging, in phenol red-free, serum-free DMEM media (Gibco, ThermoFisher Scientific) supplemented with 20 mM HEPES, with imaging at one frame per second for 5 min.

### Dual Iterative Structured Illumination Microscopy (diSIM)

#### Imaging

TMEM127-KO or Mock-KO cells stably expressing EGFP-tagged clathrin light chain were grown on #1.5 coverslips. Cells were fixed in 4% paraformaldehyde and mounted in Prolong Glass mounting medium (ThermoFisher, Waltham, MA). Cells were imaged with a Plan-Apochromat 63x/1.4 Oil immersion objective on an ELYRA7 microscope in lattice SIM mode (Carl Zeiss, White Plains, NY). A 488 nm laser was passed through a 23 µm grating and used to excite EGFP.

Fluorescence emission was collected on a sCMOS camera (PCO, Kelheim, Germany) through a 495-550 nm bandpass emission filter. 15 illumination phases were collected with an exposure time of 100 ms each. Voxel dimensions were 0.063 nm, 0.063 nm, and 0.110 nm (X, Y, Z), respectively. Z-stacks of approximately 750 nm were acquired for each field of view.

#### Processing

3D diSIM processing was performed in ZEN Black 3.0 SR edition (Carl Zeiss White Plains, NY) using the “Standard Fixed” setting. For initial visual inspections, maximum intensity projections of ∼740 nm Z-stacks were performed in ImageJ/Fiji. Z-stacks encompassing entire cells were segmented using the “Blobs Finder” tool in Vision4D (Aviris; Rostock, Germany) and rendered in 3D. For quantitative analysis of CCP diameter and number, single Z-planes in or near the lower plasma membrane were selected and run through a custom ImageJ/Fiji macro that counted CCPs and determined their 2D area.

### TIRFM Experiment Analysis

#### Fixed-cell analysis of RET Co-Localization with Clathrin Labelled Structures (CLSs)

Unbiased detection and analysis of clathrin diffraction-limited structures was performed using CME analysis software in Matlab (Mathworks Corporation, Natick, MA, RRID:SCR_001622; github.com/Danuserlab/CMEAnalysis), as previously described (Aguet et al., 2013; Cabral-Dias et al., 2022; Delos Santos et al., 2017). Briefly, diffraction-limited clathrin structures were detected using a Gaussian-based model to approximate the point-spread function of CLSs (‘primary’ channel). RET proteins (TIRFM images) or clathrin (epifluorescence images) were detected in a ‘secondary’ channel and the intensity within the CLSs, beyond local background amounts, signified the level of colocalization of the proteins.

#### Live-Cell Analysis of CCP Dynamics

Automated detection, tracking and analysis of CCPs following time-lapse TIRF microscopy of SH-SY5Y cells stably expressing EGFP-CLC was as previously described (Aguet et al., 2013; Kadlecova et al., 2017; Mettlen and Danuser, 2014). Diffraction-limited clathrin structures were detected using a Gaussian-based model method to approximate the point-spread function (Aguet et al., 2013), and trajectories were determined from clathrin structure detections using u-track (Jaqaman et al., 2008). sCLSs were distinguished from bona fide CCPs based on unbiased analysis of clathrin intensities in the early CCP stages (Aguet et al., 2013; Kadlecova et al., 2017). Both sCLSs and CCPs represent nucleation events, but only bona fide CCPs undergo stabilization, maturation and in some cases scission to produce vesicles (Aguet et al., 2013; Kadlecova et al., 2017). We report the sCLS nucleation, CCP initiation, and the fraction of CCPs <15s. Because CCPs are diffraction-limited objects, the amplitude of the Gaussian model of the fluorescence intensity of EGFP-CLC informs about CCP size (Figure 5A-E).

### Single Particle Tracking

Single particle tracking experiments were performed by labeling EGFR with a Cy3B-conjugated Fab fragment targeting the EGFR ectodomain, generated from mAb108 (Sugiyama et al., 2023). Previous characterization of this labeling strategy indicated that antibody labeling did not impact EGFR ligand binding or signaling (Sugiyama et al., 2023). Cells were labelled with 50 ng/mL Fab-Cy3B (to label and track total EGFR) for 10 min followed by washing to remove unbound antibodies, in media lacking EGF. Time-lapse TIRF imaging was performed using a Quorum (Guelph, ON, Canada) Diskovery system, comprised of a Leica DMi8 microscope with a 63×/1.49 NA TIRF objective with a 1.8× camera relay (total magnification 108×), using 561 nm laser illumination and a 620/60 nm emission filter. Images were acquired using an iXon Ultra 897 EM-CCD (Spectral Applied Research, Andor, Toronto, ON, Canada). All live cell imaging was performed using cells incubated in serum-free DMEM/F-12 or DMEM without phenol red or P/S, with a frame-rate of 20 Hz, for total length of time-lapse 250 frames. Single particles labelled with Cy3B were detected and tracked (Jaqaman et al., 2008), and the motion types and diffusion coefficients were determined using moment-scaling spectrum analysis (MSS) (Freeman et al., 2018; Jaqaman et al., 2011).

### Statistical Analyses

A minimum of three independent replicates were performed for each experiment, except where indicated. Consistency between experimental replicates was assessed through two-tailed unpaired t-tests, and One- or Two-way ANOVA with indicated multiple comparison tests using GraphPad Prism version 9 (GraphPad Prism Software, San Diego, CA, USA). Specific significance tests were selected using GraphPad Prism following residual testing for Gaussian distribution (Normality tests of Anderson-Darling, D’Agostino, Shapiro-Wilk, Kolmogorov-Smirnov) and testing whether residuals are clustered or heteroscedastic (Brown-Forsyth and Bartlett’s tests). Bar and line graphs, used for immunoblots and time course experiments, were presented with means and standard deviation (±SD), while box plots with whiskers from minimum to maximum values were used for individual cell imaging. Graphs were produced using GraphPad Prism.

## ACKNOWLEDGEMENTS

The authors would like to thank Juliana Shizas, Montdher Hussain and Kasra Ranjbar for technical assistance. This work was supported by operating grants from the Cancer Research Society of Canada (CRS-19439), Canadian Institutes for Health Research (MOP-142303, PJT-178274, LMM) and (PJT-156355, CNA), and from NIH (GM114102 and CA264248), Neuroendocrine Tumor Research Foundation, VHL Alliance, and the UT System Star Awards (PLMD). PLMD holds the Robert Tucker Hayes Distinguished Chair in Oncology. LMM is the Bracken Chair in Genetics and Molecular Medicine at Queen’s University.

## Supplementary Figures

**Figure 2-figure supplement 1.**
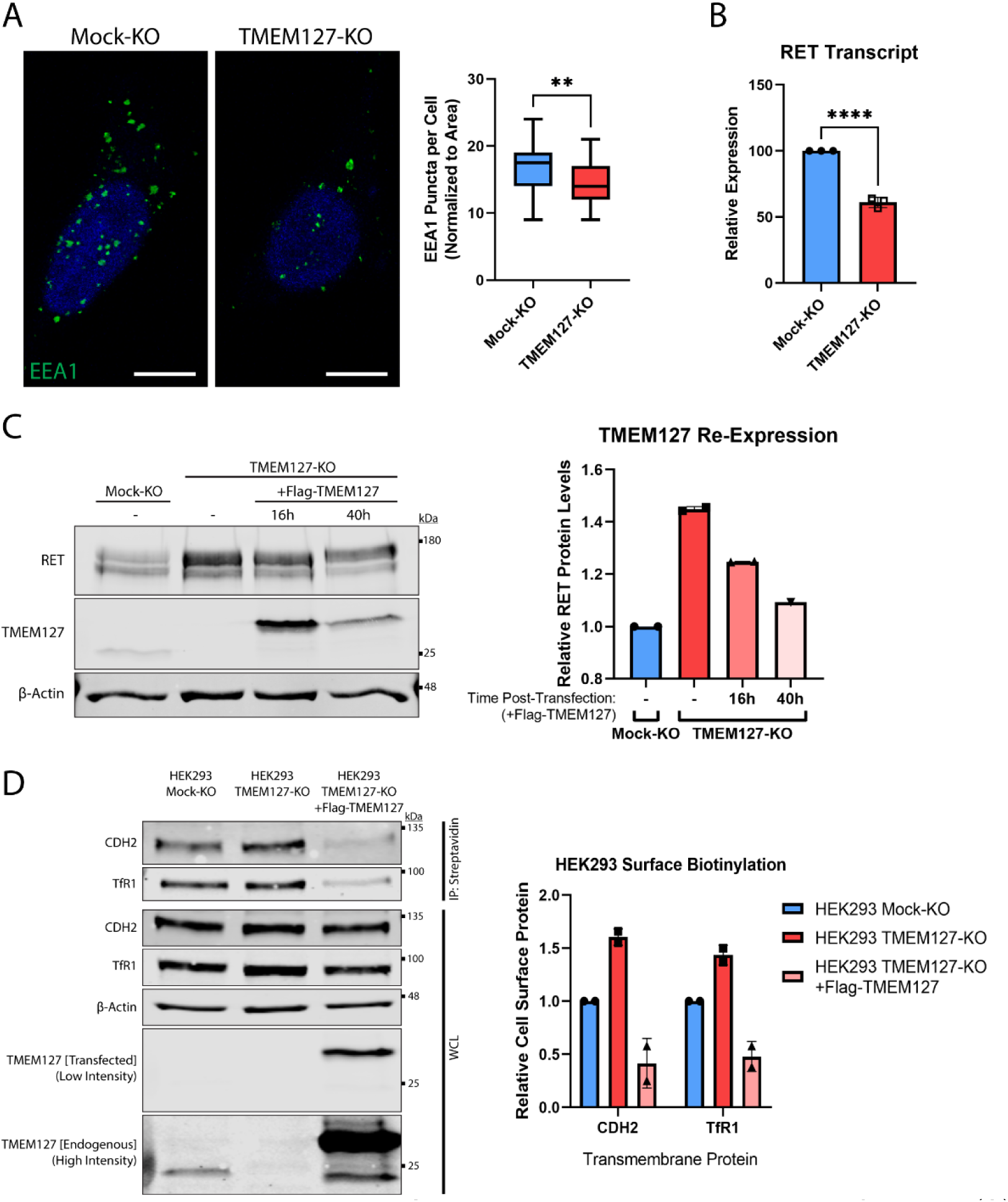
Characterization of TMEM127-KO cells. (**A**) Immunofluorescence confocal images (left panel) of Mock-KO and TMEM127-KO SH-SY5Y cells stained for early endosome marker EEA1 (green) and Hoechst nuclear stain (blue). Quantification of EEA1 puncta per cell (right panel), normalized to cell area, of n=32-43 cells per condition are shown. (Two-tailed unpaired t-test; **p<0.01). Scale bars = 5 μm. (**B**) Quantification of RET transcripts by qRT-PCR of RNA isolated from Mock-KO and TMEM127-KO SH-SY5Y cells as previously described (Richardson et al., 2009). RET transcript levels are expressed relative to Mock-KO cells. Three independent experiments (n=3) are shown as mean ±SD (Two-tailed unpaired t-test; ****p<0.0001). (**C**) Immunoblots showing RET protein expression in Mock-KO and TMEM127-KO SH-SY5Y cells (left panel) upon transient re-expression of Flag-tagged wildtype TMEM127 (+Flag-TMEM127). Period following initial transfection indicated in hours (h). Quantification of RET protein levels normalized to β-Actin loading control and expressed relative to Mock-KO (right panel). RET protein levels in TMEM127-KO decrease and approach Mock-KO cell levels with longer transient TMEM127 expression. Rescue limited by relatively low transfection efficiency in SH-SY5Y cells. **(D)** Immunoblots showing the indicated total and cell surface biotinylated transmembrane proteins in Mock-KO and in TMEM127-KO HEK293 cells with and without add back of wildtype TMEM127 (+Flag-TMEM127) (left panel). High and low intensity images of a single blot show differential levels of endogenous and transfected Flag-TMEM127. β-Actin was used as a loading control. Biotinylated proteins N-Cadherin (CDH2), and Transferrin Receptor-1 (TfR1) were collected, separated, and immunoblotted as in Figure 2A. Quantification of biotinylated cell surface protein levels (IP) normalized to corresponding total protein (WCL) and expressed relative to Mock-KO for each protein is shown (right panel).

**Figure 2-figure supplement 2.**
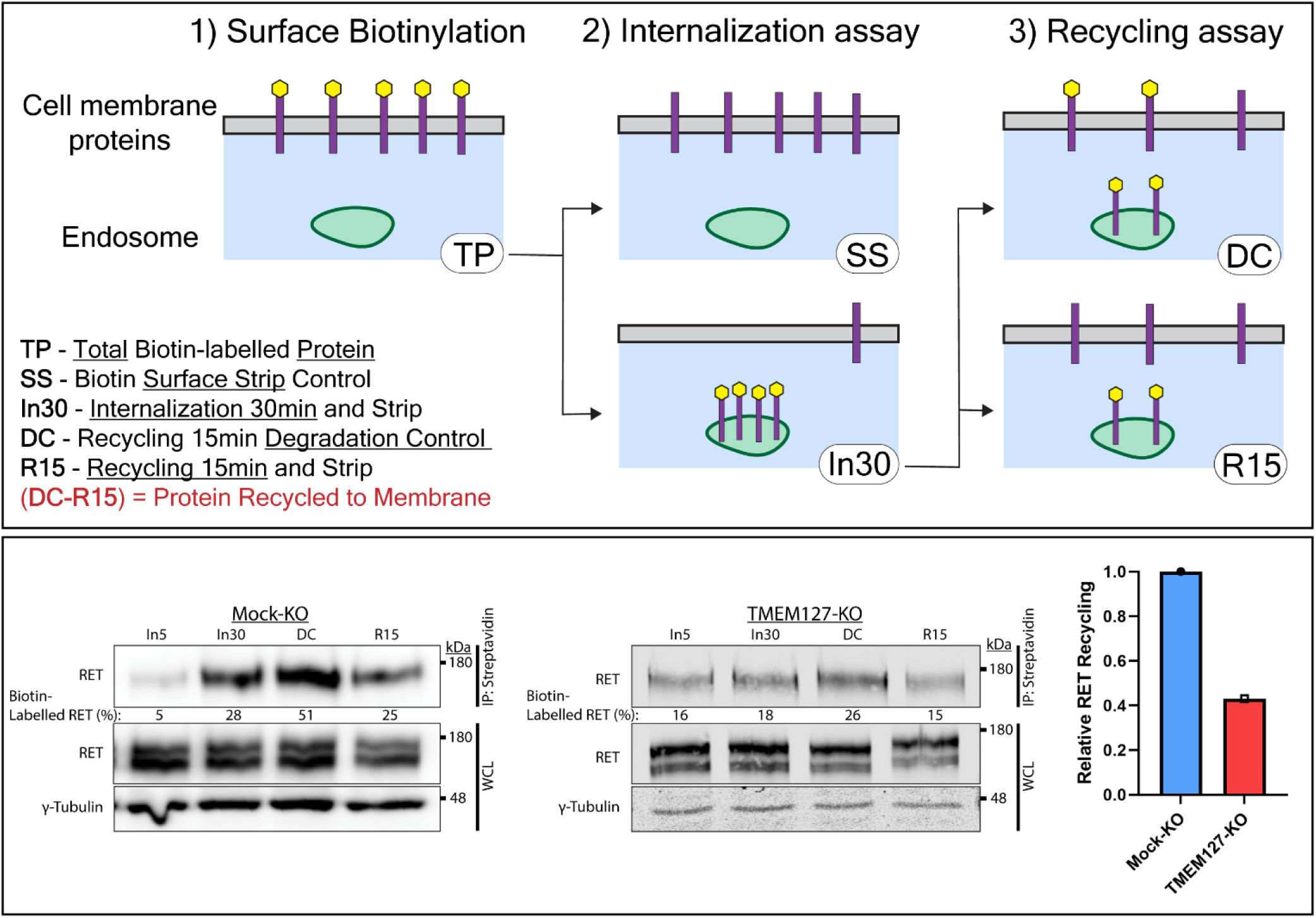
RET recycling is reduced in the absence of TMEM127. The recycling assay is an extension of the internalization assay shown in Figure 2B, where an additional incubation allows for receptor recycling back to the cell membrane. (**upper panel**) Diagram of biotin assay workflow. Surface proteins are biotin labelled (TP; total protein) and either stripped of cell-surface biotin immediately (SS; Surface Strip control) or incubated at 37°C with ligand for 30 min (In30) to allow internalization, and remaining cell-surface biotin is stripped. To evaluate recycling, In30 cells undergo an additional 15 min incubation to allow protein recycling to the membrane (DC; Degradation Control) followed by another biotin stripping step (R15; Recycling 15 min). The portion of RET receptors recycled is represented by the difference in intensity between DC and R15. (**lower panel**) Immunoblots showing recycling of biotinylated cell-surface RET protein in Mock-KO (left) and TMEM127-KO (middle) SH-SY5Y cells. Biotinylated proteins were collected, separated, and immunoblotted as in Figure 2A. γ-Tubulin was used as a loading control. Quantification of RET Recycling (right) shown relative to Mock-KO cells.

**Figure 8-figure supplement 1.**
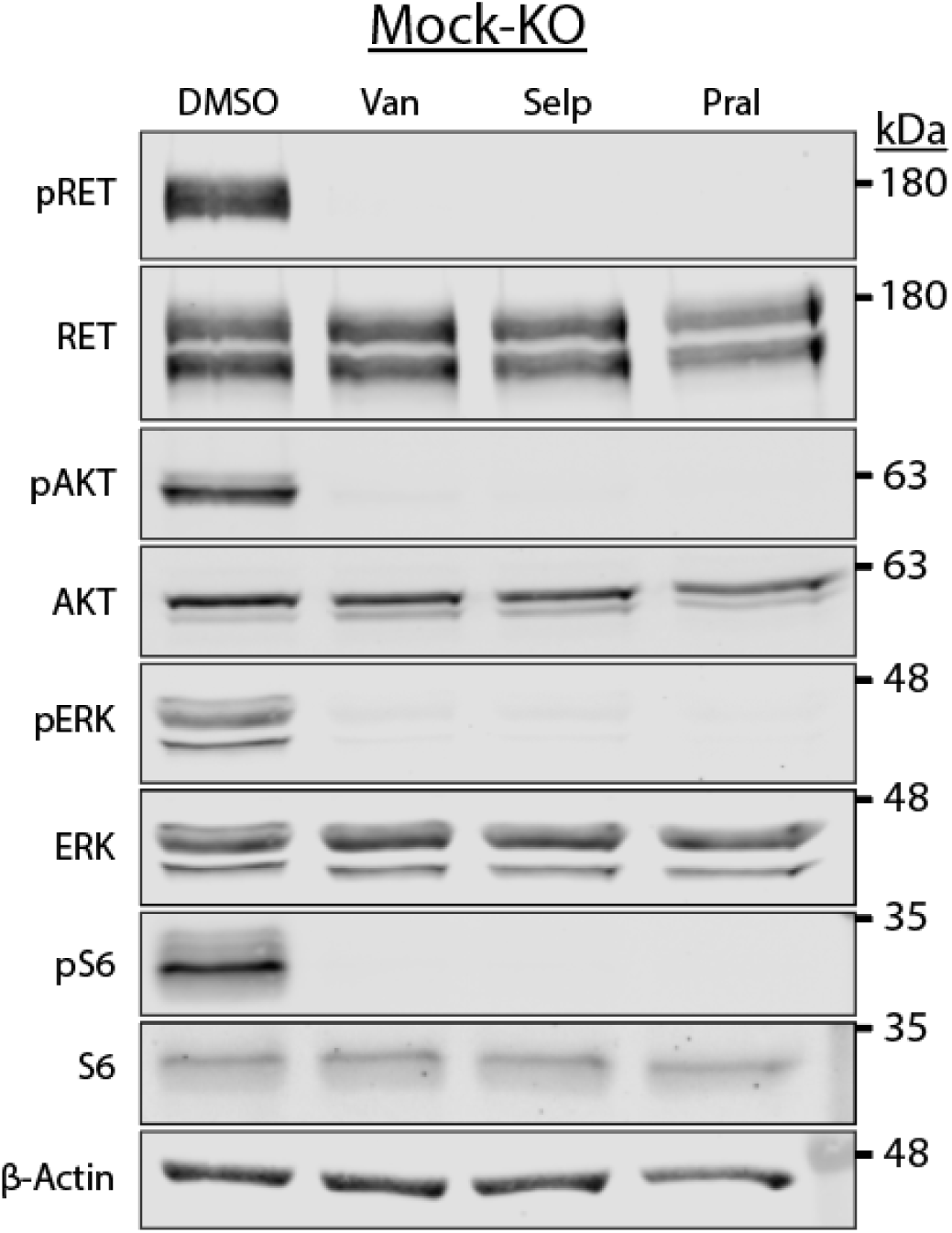
RET activation promoted AKT, ERK, and S6 phosphorylation in SH-SY5Y cells. Mock-KO SH-SY5Y cells treated with DMSO, or 5 μM Vandetanib (Van), Selpercatinib (Selp), or Pralsetinib (Pral) (1h) and then GDNF (30 min). RET, AKT, ERK and S6 phosphorylation was abrogated by all three inhibitors.

**Figure 8-figure supplement 2.**
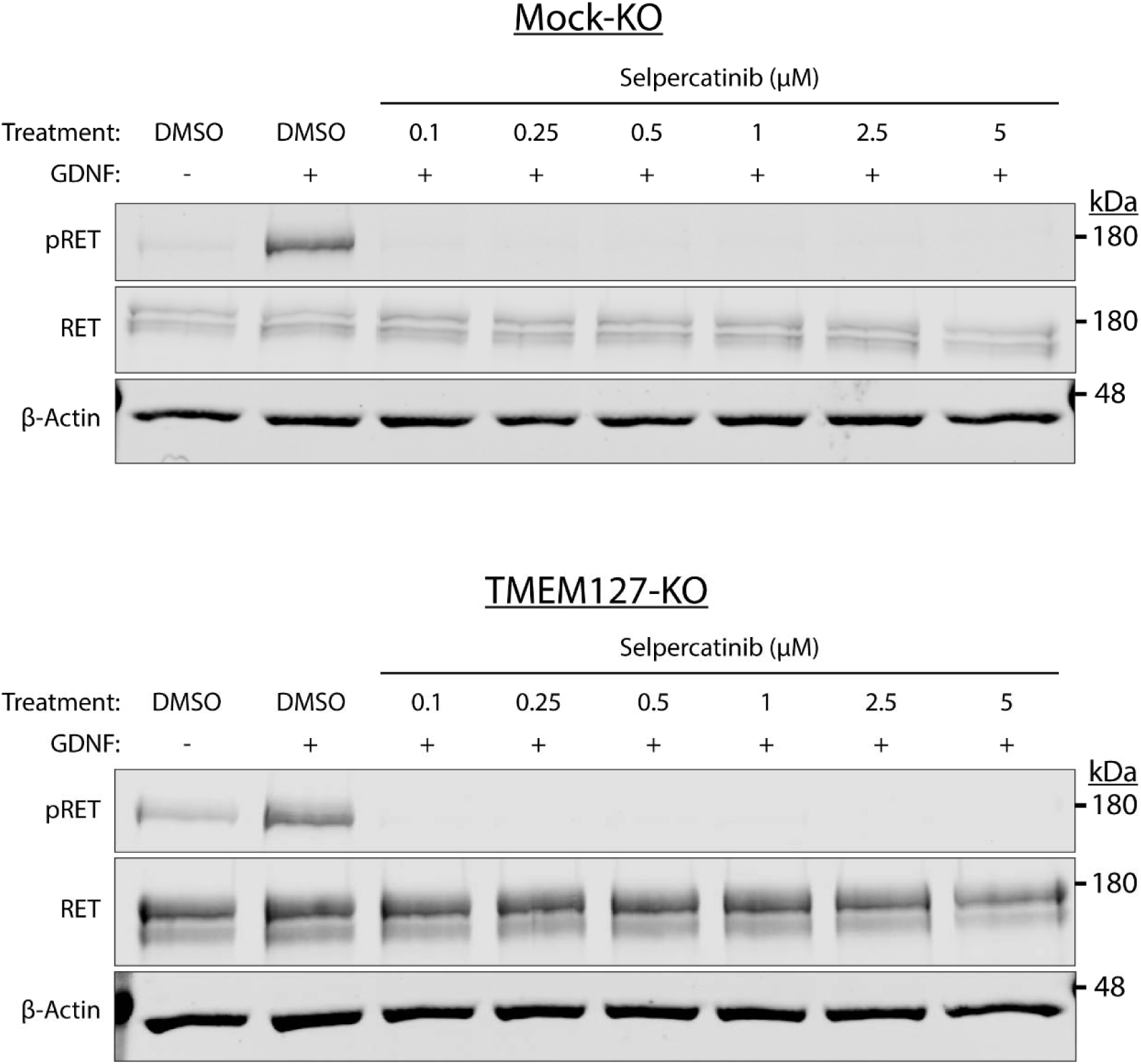
Optimization of Selpercatinib concentration. Mock-KO and TMEM127-KO SH-SY5Y cells treated with GDNF, and DMSO or indicated concentrations of Selpercatinib for 1h. RET phosphorylation was abrogated at all concentrations tested. The lowest concentration was chosen for MTT assays (Figure 8E) to inhibit RET, while avoiding cell death due to high drug concentrations.

**Figure 8-figure supplement 3.**
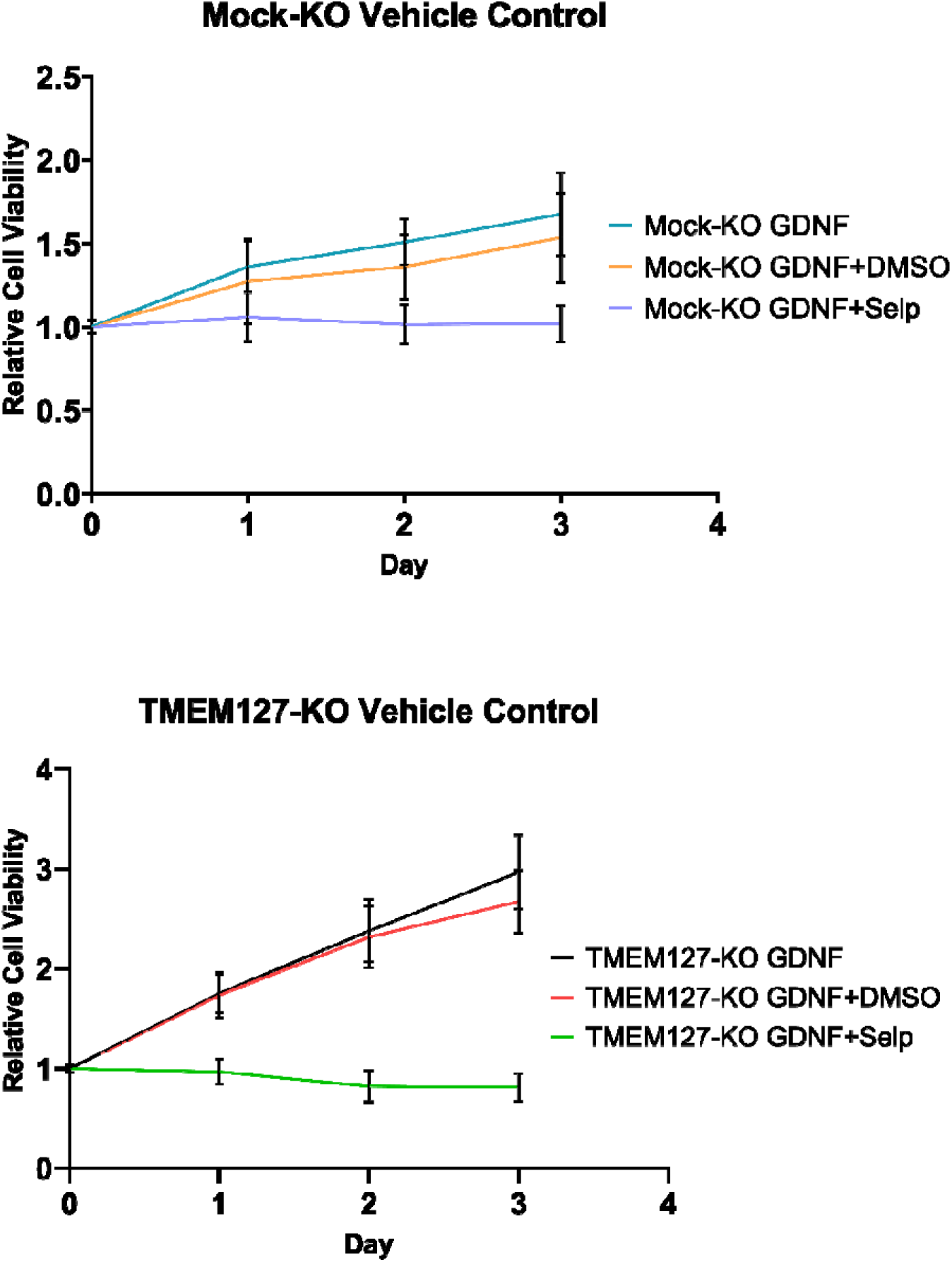
Vehicle treatment does not significantly affect SH-SY5Y cell proliferation. MTT assay of Mock-KO and TMEM127-KO SH-SY5Y cell proliferation in the presence of GDNF, and DMSO or Selpercatinib (Selp) for 1 h. The mean ±SD of three independent experiments, for each graph, with 18 replicates per condition (n=54) are shown.

## Supplementary Animation

**Animation 1.**
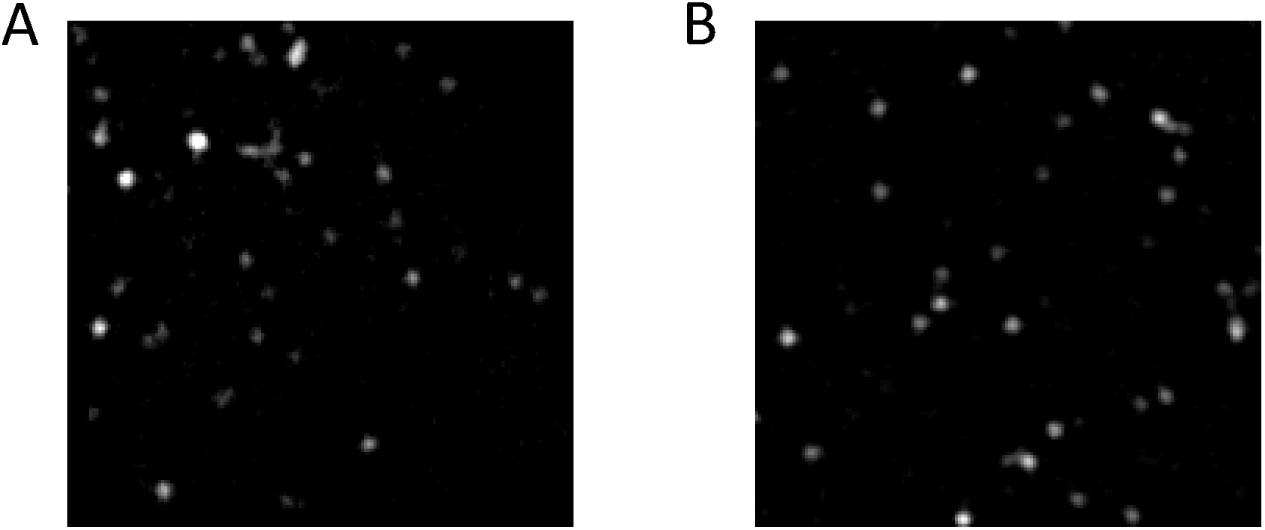
Assembly of clathrin-labelled structures is reduced in the absence of TMEM127. – Live cell TIRFM imaging movies of cell surface clathrin in Mock-KO and TMEM127-KO SH-SY5Y cells expressing EGFP-tagged clathrin light chain. Timelapse videos of cells (5 min @ 1 frame-per-second) are shown. (**A**) In Mock-KO cells clathrin-labelled structures (CLS) assemble rapidly and abundantly. These may abort (very short-lived puncta) or progress to mature CCP before internalizing (∼40 seconds) and disappearing from the TIRF field (Crupi et al., 2015). (**B**) TMEM127-KO cells show fewer CLS, which assemble more slowly and remain on the membrane much longer before moving out of the TIRF field, suggesting reduced recruitment of clathrin to CLS in these cells, contributing to reduced ability of CCPs to mature and capture protein cargo.

## REFERENCES

Aguet, F., C.N. Antonescu, M. Mettlen, S.L. Schmid, and G. Danuser. 2013. Advances in analysis of low signal-to-noise images link dynamin and AP2 to the functions of an endocytic checkpoint. Dev Cell. 26:279–291.

Cabral-Dias, R., S. Lucarelli, K. Zak, S. Rahmani, G. Judge, J. Abousawan, L.F. DiGiovanni, D. Vural, K.E. Anderson, M.G. Sugiyama, G. Genc, W. Hong, R.J. Botelho, G.D. Fairn, P.K. Kim, and C.N. Antonescu. 2022. Fyn and TOM1L1 are recruited to clathrin-coated pits and regulate Akt signaling. The Journal of cell biology. 221.

Chen, X., Q. Lu, H. Zhou, J. Liu, B. Nadorp, A. Lasry, Z. Sun, B. Lai, G. Rona, J. Zhang, M. Cammer, K. Wang, W. Al-Santli, Z. Ciantra, Q. Guo, J. You, D. Sengupta, A. Boukhris, H. Zhang, C. Liu, P. Cresswell, P.L.M. Dahia, M. Pagano, I. Aifantis, and J. Wang. 2023. A membrane-associated MHC-I inhibitory axis for cancer immune evasion. Cell. 186:3903–3920 e3921.

Crupi, M.J.F., S.M. Maritan, E. Reyes-Alvarez, E.Y. Lian, B.D. Hyndman, A.N. Rekab, S. Moodley, C.N. Antonescu, and L.M. Mulligan. 2020. GGA3-mediated recycling of the RET receptor tyrosine kinase contributes to cell migration and invasion. Oncogene. 39:1361–1377.

Crupi, M.J.F., P. Yoganathan, L.N. Bone, E. Lian, A. Fetz, C.N. Antonescu, and L.M. Mulligan. 2015. Distinct Temporal Regulation of RET Isoform Internalization: Roles of Clathrin and AP2. Traffic. 16:1155–1173.

Dahia, P.L. 2014. Pheochromocytoma and paraganglioma pathogenesis: learning from genetic heterogeneity. Nature reviews. Cancer. 14:108–119.

Delos Santos, R.C., S. Bautista, S. Lucarelli, L.N. Bone, R.M. Dayam, J. Abousawan, R.J. Botelho, and C.N. Antonescu. 2017. Selective regulation of clathrin-mediated epidermal growth factor receptor signaling and endocytosis by phospholipase C and calcium. Mol Biol Cell. 28:2802–2818.

Deng, Y., Y. Qin, S. Srikantan, A. Luo, Z.M. Cheng, S.K. Flores, K.S. Vogel, E. Wang, and P.L.M. Dahia. 2018. The TMEM127 human tumor suppressor is a component of the mTORC1 lysosomal nutrient-sensing complex. Hum Mol Genet. 27:1794–1808.

Djakbarova, U., Y. Madraki, E.T. Chan, and C. Kural. 2021. Dynamic interplay between cell membrane tension and clathrin-mediated endocytosis. Biol Cell. 113:344–373.

Du, Z., and C.M. Lovly. 2018. Mechanisms of receptor tyrosine kinase activation in cancer. Mol Cancer. 17:58.

Ehrlich, M., W. Boll, A. Van Oijen, R. Hariharan, K. Chandran, M.L. Nibert, and T. Kirchhausen. 2004. Endocytosis by random initiation and stabilization of clathrin-coated pits. Cell. 118:591–605.

Flores, S.K., Y. Deng, Z. Cheng, X. Zhang, S. Tao, A. Saliba, I. Chu, N. Burnichon, A.P. Gimenez-Roqueplo, E. Wang, R.C.T. Aguiar, and P.L.M. Dahia. 2020. Functional Characterization of TMEM127 Variants Reveals Novel Insights into Its Membrane Topology and Trafficking. J Clin Endocrinol Metab. 105:e3142–3156.

Freeman, S.A., A. Vega, M. Riedl, R.F. Collins, P.P. Ostrowski, E.C. Woods, C.R. Bertozzi, M.I. Tammi, D.S. Lidke, P. Johnson, S. Mayor, K. Jaqaman, and S. Grinstein. 2018. Transmembrane Pickets Connect Cyto- and Pericellular Skeletons Forming Barriers to Receptor Engagement. Cell. 172:305–317 e310.

Guo, Q., Z.M. Cheng, H. Gonzalez-Cantu, M. Rotondi, G. Huelgas-Morales, P. Ethiraj, Z. Qiu, J. Lefkowitz, W. Song, B.N. Landry, H. Lopez, C.M. Estrada-Zuniga, S. Goyal, M.A. Khan, T.J. Walker, E. Wang, F. Li, Y. Ding, L.M. Mulligan, R.C.T. Aguiar, and P.L.M. Dahia. 2023. TMEM127 suppresses tumor development by promoting RET ubiquitination, positioning, and degradation. Cell Rep. 42:113070.

Horton, C., H. LaDuca, A. Deckman, K. Durda, M. Jackson, M.E. Richardson, Y. Tian, A. Yussuf, K. Jasperson, and T. Else. 2022. Universal Germline Panel Testing for Individuals With Pheochromocytoma and Paraganglioma Produces High Diagnostic Yield. J Clin Endocrinol Metab. 107:e1917–e1923.

Hyndman, B.D., M.J.F. Crupi, S. Peng, L.N. Bone, A.N. Rekab, E.Y. Lian, S.M. Wagner, C.N. Antonescu, and L.M. Mulligan. 2017. Differential recruitment of E3 ubiquitin ligase complexes regulates RET isoform internalization. J Cell Sci. 130:3282–3296.

Jaqaman, K., H. Kuwata, N. Touret, R. Collins, W.S. Trimble, G. Danuser, and S. Grinstein. 2011. Cytoskeletal control of CD36 diffusion promotes its receptor and signaling function. Cell. 146:593–606.

Jaqaman, K., D. Loerke, M. Mettlen, H. Kuwata, S. Grinstein, S.L. Schmid, and G. Danuser. 2008. Robust single-particle tracking in live-cell time-lapse sequences. Nat Methods. 5:695–702.

Kadlecova, Z., S.J. Spielman, D. Loerke, A. Mohanakrishnan, D.K. Reed, and S.L. Schmid. 2017. Regulation of clathrin-mediated endocytosis by hierarchical allosteric activation of AP2. The Journal of cell biology. 216:167–179.

Kaksonen, M., and A. Roux. 2018. Mechanisms of clathrin-mediated endocytosis. Nat Rev Mol Cell Biol. 19:313–326.

Le Hir, H., L.G. Colucci-D’Amato, N. Charlet-Berguerand, P.F. Plouin, X. Bertagna, V. de Franciscis, and C. Thermes. 2000. High levels of tyrosine phosphorylated proto-ret in sporadic phenochromocytomas. Cancer Res. 60:1365–1370.

Löschberger, A., Y. Novikau, R. Netz, M.-C. Spindler, R. Benavente, T. Klein, M. Sauer, and D.I. Kleppe. 2021. Super-Resolution Imaging by Dual Iterative Structured Illumination Microscopy. bioRxiv:2021.2005.2012.443720.

Mettlen, M., P.H. Chen, S. Srinivasan, G. Danuser, and S.L. Schmid. 2018. Regulation of Clathrin-Mediated Endocytosis. Annu Rev Biochem. 87:871–896.

Mettlen, M., and G. Danuser. 2014. Imaging and modeling the dynamics of clathrin-mediated endocytosis. Cold Spring Harb Perspect Biol. 6:a017038.

Mulligan, L.M. 2019. GDNF and the RET Receptor in Cancer: New Insights and Therapeutic Potential. Frontiers in Physiology. 9:1873.

Neumann, H.P.H., W.F. Young, Jr., and C. Eng. 2019. Pheochromocytoma and Paraganglioma. N Engl J Med. 381:552–565.

Pierchala, B.A., J. Milbrandt, and E.M. Johnson, Jr. 2006. Glial cell line-derived neurotrophic factor-dependent recruitment of Ret into lipid rafts enhances signaling by partitioning Ret from proteasome-dependent degradation. J Neurosci. 26:2777–2787.

Qin, Y., Y. Deng, C.J. Ricketts, S. Srikantan, E. Wang, E.R. Maher, and P.L. Dahia. 2014. The tumor susceptibility gene TMEM127 is mutated in renal cell carcinomas and modulates endolysosomal function. Hum Mol Genet. 23:2428–2439.

Qin, Y., L. Yao, E.E. King, K. Buddavarapu, R.E. Lenci, E.S. Chocron, J.D. Lechleiter, M. Sass, N. Aronin, F. Schiavi, F. Boaretto, G. Opocher, R.A. Toledo, S.P. Toledo, C. Stiles, R.C. Aguiar, and P.L. Dahia. 2010. Germline mutations in TMEM127 confer susceptibility to pheochromocytoma. Nat Genet. 42:229–233.

Reyes-Alvarez, E., T.J. Walker, and L.M. Mulligan. 2022. Evaluating Cell Membrane Localization and Intracellular Transport of Proteins by Biotinylation. Methods Mol Biol. 2508:197–209.

Richardson, D.S., T.S. Gujral, S. Peng, S.L. Asa, and L.M. Mulligan. 2009. Transcript level modulates the inherent oncogenicity of RET/PTC oncoproteins. Cancer Res. 69:4861–4869.

Richardson, D.S., A.Z. Lai, and L.M. Mulligan. 2006. RET ligand-induced internalization and its consequences for downstream signaling. Oncogene. 25:3206–3211.

Richardson, D.S., D.M. Rodrigues, B.D. Hyndman, M.J. Crupi, A.C. Nicolescu, and L.M. Mulligan. 2012. Alternative splicing results in RET isoforms with distinct trafficking properties. Mol Biol Cell. 23:3838–3850.

Saleem, M., S. Morlot, A. Hohendahl, J. Manzi, M. Lenz, and A. Roux. 2015. A balance between membrane elasticity and polymerization energy sets the shape of spherical clathrin coats. Nature communications. 6:6249.

Schmid, S.L. 2017. Reciprocal regulation of signaling and endocytosis: Implications for the evolving cancer cell. The Journal of cell biology. 216:2623–2632.

Sonnino, S., L. Mauri, V. Chigorno, and A. Prinetti. 2007. Gangliosides as components of lipid membrane domains. Glycobiology. 17:1R–13R.

Sorkin, A., and A. Fortian. 2015. Endocytosis and Endosomal Sorting of Receptor Tyrosine Kinases. In Receptor Tyrosine Kinases: Structure, Functions and Role in Human Disease. D. Wheeler and Y. Yarden, editors. Springer, New York, NY.

Sugiyama, M.G., A.I. Brown, J. Vega-Lugo, J.P. Borges, A.M. Scott, K. Jaqaman, G.D. Fairn, and C.N. Antonescu. 2023. Confinement of unliganded EGFR by tetraspanin nanodomains gates EGFR ligand binding and signaling. Nature communications. 14:2681.

Takahashi, M., Y. Buma, and M. Taniguchi. 1991. Identification of the *ret* proto-oncogene products in neuroblastoma and leukemia cells. Oncogene. 6:297–301.

Takaya, K., T. Yoshimasa, H. Arai, N. Tamura, Y. Miyamoto, H. Itoh, and K. Nakao. 1996a. Expression of the RET proto-oncogene in normal human tissues, pheochromocytomas, and other tumors of neural crest origin. J Mol Med. 74:617–621.

Takaya, K., T. Yoshimasa, H. Arai, N. Tamura, Y. Miyamoto, H. Itoh, and K. Nakao. 1996b. The RET proto-oncogene in sporadic pheochromocytomas. Intern Med. 35:449–452.

Tate, J.G., S. Bamford, H.C. Jubb, Z. Sondka, D.M. Beare, N. Bindal, H. Boutselakis, C.G. Cole, C. Creatore, E. Dawson, P. Fish, B. Harsha, C. Hathaway, S.C. Jupe, C.Y. Kok, K. Noble, L. Ponting, C.C. Ramshaw, C.E. Rye, H.E. Speedy, R. Stefancsik, S.L. Thompson, S. Wang, S. Ward, P.J. Campbell, and S.A. Forbes. 2019. COSMIC: the Catalogue Of Somatic Mutations In Cancer. Nucleic Acids Res. 47:D941–D947.

Toledo, R.A., N. Burnichon, A. Cascon, D.E. Benn, J.P. Bayley, J. Welander, C.M. Tops, H. Firth, T. Dwight, T. Ercolino, M. Mannelli, G. Opocher, R. Clifton-Bligh, O. Gimm, E.R. Maher, M. Robledo, A.P. Gimenez-Roqueplo, P.L. Dahia, and N.S. Group. 2017. Consensus Statement on next-generation-sequencing-based diagnostic testing of hereditary phaeochromocytomas and paragangliomas. Nat Rev Endocrinol. 13:233–247.

von Zastrow, M., and A. Sorkin. 2021. Mechanisms for Regulating and Organizing Receptor Signaling by Endocytosis. Annu Rev Biochem. 90:709–737.

Wu, X., X. Zhao, L. Baylor, S. Kaushal, E. Eisenberg, and L.E. Greene. 2001. Clathrin exchange during clathrin-mediated endocytosis. The Journal of cell biology. 155:291–300.

